# Contrasting genome dynamics between domesticated and wild yeasts

**DOI:** 10.1101/076562

**Authors:** Jia-Xing Yue, Jing Li, Louise Aigrain, Johan Hallin, Karl Persson, Karen Oliver, Anders Bergström, Paul Coupland, Jonas Warringer, Marco Consentino Lagomarsino, Gilles Fischer, Richard Durbin, Gianni Liti

## Abstract

Structural rearrangements have long been recognized as an important source of genetic variation with implications in phenotypic diversity and disease, yet their evolutionary dynamics are difficult to characterize with short-read sequencing. Here, we report long-read sequencing for 12 strains representing major subpopulations of the partially domesticated yeast *Saccharomyces cerevisiae* and its wild relative *Saccharomyces paradoxus*. Complete genome assemblies and annotations generate population-level reference genomes and allow for the first explicit definition of chromosome partitioning into cores, subtelomeres and chromosome-ends. High-resolution view of structural dynamics uncovers that, in chromosomal cores, *S. paradoxus* exhibits higher accumulation rate of balanced structural rearrangements (inversions, translocations and transpositions) whereas *S. cerevisiae* accumulates unbalanced rearrangements (large insertions, deletions and duplications) more rapidly. In subtelomeres, recurrent interchromosomal reshuffling was found in both species, with higher rate in *S. cerevisiae*. Such striking contrasts between wild and domesticated yeasts reveal the influence of human activities on structural genome evolution.

## Introduction

Understanding how genetic variation translates into phenotypic diversity is a central theme in genomic studies. With the rapid advancement of sequencing technology in recent decades, genetic variation in large natural populations has been extensively explored for human (The 1000 Genomes Project Consortium 2010) and several important model organisms such as yeast (Liti et al. 2009a; Bergström et al. 2014; Strope et al. 2015; Gallone et al. 2016), fruitfly (Mackay et al. 2012; Huang et al. 2014) and *Arabidopsis* (Cao et al. 2011; The 1001 Genomes Consortium 2016). However, our current knowledge of natural genetic variation is heavily biased towards single nucleotide variants (SNVs), although it is increasingly appreciated that structural variants (SVs) (i.e. inversions, translocations, and large insertions, deletions and duplications) might contribute more to the overall genetic differences and have far more significant evolutionary and functional consequences (Feuk et al. 2006; Weischenfeldt et al. 2013). For example, inversion and translocation can facilitate adaptation and speciation by suppressing local recombination in heterozygotes, which creates genetic barrier to gene flow and promotes fast differentiation between subpopulations (Rieseberg 2001). Large insertion, deletion and duplication, on the other hand, can sometimes cause diseases by altering the dosage of gene products (Weischenfeldt et al. 2013). Therefore, population and comparative studies on the landscape and dynamics of structural rearrangements are needed to understand the evolutionary balance between genome stability and plasticity as well as its functional implications in adaptation and disease.

The major challenge of studying structural rearrangements at the genomic level lies in the limited power and resolution of detecting rearrangement variants with traditional capillary and short-read sequencing data. Moreover, many structural rearrangement are embedded in complex genomic regions that are repetitive and highly dynamic, which further exacerbate this problem. For example, subtelomeric regions are known hotspots of rampant interchromosomal reshuffling which significantly contribute to inter-individual genetic variation and phenotypic diversity of eukaryotic organisms (Pryde et al. 1997; Mefford and Trask 2002; Eichler and Sankoff 2003; Dujon 2010). However, the detailed landscape and evolutionary dynamics of such subtelomeric reshuffling has remained elusive due to the difficulty in assembling these regions. The newly available long-read sequencing technology represented by PacBio and Oxford Nanopore can generate very long reads (>10 kb) with high throughput, thereby constituting powerful tools for detecting and characterizing complex structural rearrangements (Goodwin et al. 2016). Recent applications of long-read sequencing technology in several reference genome sequencing projects has proved to be quite successful, producing highly continuous genome assemblies with most of complex regions correctly resolved, even for large mammalian genomes (Chaisson et al. 2014; VanBuren et al. 2015; Gordon et al. 2016).

The baker’s yeast *S. cerevisiae* has long been used as an important model system in biological studies, illuminating almost every aspect of molecular biology and genetics. Its genome was the first to be fully sequenced and completely assembled in eukaryotes (Goffeau et al. 1996). However, it was only recently that the rich genetic variation and phenotypic diversity in natural yeast populations began to be appreciated (Liti 2015). Our first population genomics study on the partially domesticated *S. cerevisiae* and its closest wild relative *S. paradoxus* uncovered strong population differentiation in both species, which is well-correlated with their geographic origins and phenotypic diversity (Liti et al. 2009a; Warringer et al. 2011). In a following study based on high-coverage short-read sequencing, we found a surprising enrichment of genome content variation (i.e. the presence/absence of genetic materials) in the *S. cerevisiae* population despite much lower levels of SNVs as compared to the *S. paradoxus* population (Bergström et al. 2014). Here, we applied PacBio sequencing to 12 representative strains of both species, with the aim of generating a high-resolution view of the landscape and evolutionary dynamics of structural rearrangements in their genome evolution. To our knowledge, this is the first study in eukaryotes that goes beyond the scope of single reference genome sequencing and brings the PacBio sequencing technology to the population level. We generated high quality *de novo* assemblies for both nuclear and mitochondrial genomes with exceptional continuity and completeness. We further partitioned nuclear chromosomes into cores, subtelomeres and chromosome-ends to assess the structural dynamics within each partition separately. The comparison of these complete genomes coupled with explicit genome partitioning allowed us to generate a comprehensive view of structural dynamics for these two closely related species with unprecedented resolution. Our analysis highlights the influence of human activity on shaping structural genome evolution. Moreover, we report several non-canonical structures of chromosome-ends as well as lineage-specific structural rearrangements and introgression in mitochondrial genomes. In addition, we used two case studies of complex multi-allelic loci to illustrate how the precisely characterized structural rearrangements based on our complete genome assembly and annotation can be further connected to phenotypic diversity. Finally, we believe our collection of high quality annotated assemblies can serve as alternative reference genomes to guide future genomic and functional studies in yeasts.

## Results

### Complete genome assemblies across the *S. cerevisiae* and *S. paradoxus* subpopulations provide new population-level reference genomes

We selected seven *S. cerevisiae* and five *S. paradoxus* strains (Table S1) to represent previously identified evolutionary distinct subpopulations of these two species (Liti et al. 2009a; Bergström et al. 2014). For each strain, we sequenced its haploid (or homozygous diploid) genome using deep PacBio (the P6-C4 chemistry) (100-300x) and Illumina (200-500x) sequencing (Table S2). The raw PacBio *de novo* assemblies of both nuclear and mitochondrial genomes exhibited compelling quality in terms of both completeness and accuracy. Most chromosomes were assembled into single contigs with telomeric ends correctly assembled. We compared our assembly of the *S. cerevisiae* reference strain S288c with previously reported assemblies based on various sequencing technologies (PacBio, Oxford Nanopore, and Illumina MiSeq) (Kim et al. 2014; Goodwin et al. 2015). Our PacBio assembly outperformed other assemblies in terms of completeness, especially for the challenging genomic regions (e.g. Ty retrotransposable elements, telomeres and subtelomeres) (Figure S1). To further boost the assembly quality, we performed manual gap filling by referring to assemblies generated for the same strain in the early phase of this project using the older PacBio chemistry (P4-C2). We also carried out error correction based on the Illumina read alignments to minimize the remaining sequencing errors (Table S3 and S4). For each final assembly, we conducted comprehensive annotation for various genomic features, including centromeres, protein-coding genes, tRNAs, Ty retrotransposable elements, core X-elements, Y'-elements and mitochondrial RNAs (Table S5-S7).

Our final genome assemblies show outstanding completeness and accuracy compared with the current *S. cerevisiae* and *S. paradoxus* reference genomes. In general, the genome-wide dotplot comparison between these two reference genomes and our PacBio assemblies of the same strains (S288c for *S. cerevisiae* and CBS432 for *S. paradoxus*) revealed clean colinearity for both nuclear and mitochondrial genomes, suggesting comparable quality at this resolution (Figure 1A and 1B). While we noticed a few discrepancies when zooming into individual chromosome, further analyses suggested that most, if not all, of these discrepancies were due to assembly problems in the current reference genomes. For example, we found five *bona-fide* Ty1 insertions on S288c chrIII in our assembly but not in the current *S. cerevisiae* reference genome (Figure 1A, inset). These Ty1 insertions were further confirmed both by previous studies (Wheelan et al. 2006; Shibata et al. 2009; Hoang et al. 2010) and by our own long-range PCR amplifications. In the original *S. cerevisiae* reference genome sequencing project, chrIII was sequenced from several closely related but not identical strains (Oliver et al. 1992), which might explain the inconsistency. Likewise, we found a clear mis-assembly on chrIV (Figure 1B, inset) in the current *S. paradoxus* reference genome for the strain CBS432, which is confirmed both by the cross-comparison among different *S. paradoxus* strains and by the read mapping using Illumina reads and previously generated Sanger reads (Liti et al. 2009a). Moreover, we checked a few known cases of copy number variation (CNV) (e.g. Y’-elements (Liti et al. 2005), the *CUP1* (Bergström et al. 2014) and *ARR* (Bergström et al. 2014) gene clusters) and structural rearrangements (e.g. those in the Malaysian *S. cerevisiae* UWOPS03-461.4 (Marie-Nelly et al. 2014)) and they were all correctly recaptured in our assemblies, which further proves the quality of our assemblies.

**Figure 1.**
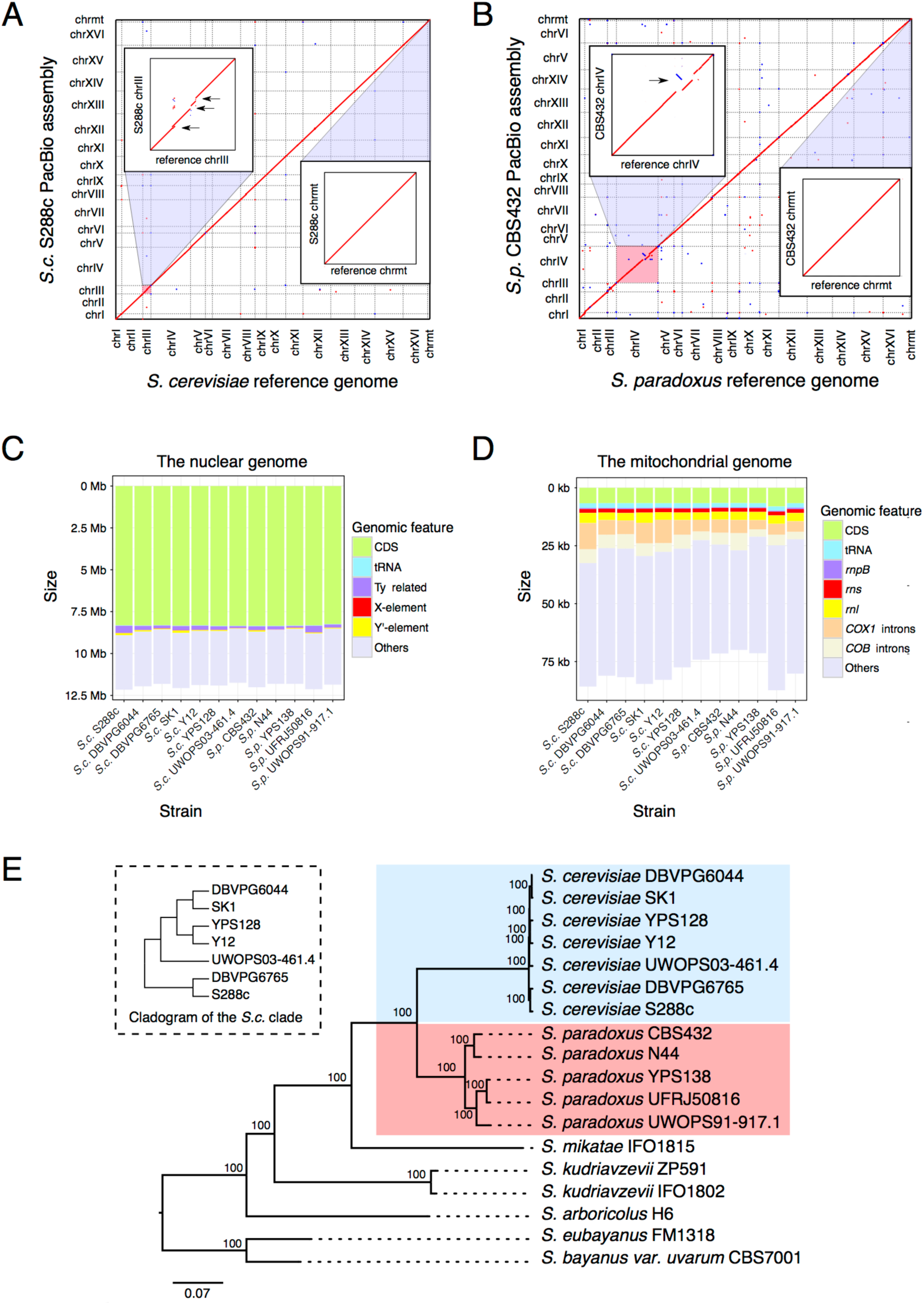
Complete genome assemblies and phylogenetic framework. (A) Dotplot comparison between the *S. cerevisiae* reference genome (strain: S288c; version: SGD R64-1-1) (X-axis) and our S288c PacBio assembly (Y-axis). Sequence homology signals were depicted in red (forward match) or blue (reverse match). The two insets show the zoomed-in plot for chromosome III (chrIII) and the mitochondrial genome (chrmt) respectively. The three black arrows indicate the Ty-containing regions (containing 5 full-length Ty1) missing in the *S. cerevisiae* reference assembly. (B) Dotplot for the comparison between the *S. paradoxus* reference genome (strain: CBS432) (X-axis) and our CBS432 PacBio assembly (Y-axis), color coded as in panel A. The two insets show the zoomed-in plot for chromosome 04 (chr04) and the mitochondrial genome (chrmt) respectively. The black arrow indicates the misassembly on chr04 in the *S. paradoxus* reference genome. (C-D) Cumulative sequence length of different annotated genomic features relative to the overall size of the *de novo* assembled nuclear genomes (panel C) and mitochondrial genomes (panel D). (E) The phylogenetic relationship of the seven *S. cerevisiae* strains (highlighted in light blue) and five *S. paradoxus* strains (highlighted in light red) sequenced in this study. Six strains from other closely related *Saccharomyces sensu stricto* species were used as outgroups. The maximum likelihood (ML) tree is based on the concatenated protein sequence matrix of 4,531 one-to-one orthologs across these 18 strains. All the internal nodes have 100% fast-bootstrap support. The inset shows the detailed relationship of the seven *S. cerevisiae* strains using cladogram.

The final assembly sizes of these 12 strains ranged from 11.75 to 12.16 Mb for the nuclear genome (Figure 1C and Table S8) and from 69.95 kb to 87.37 kb for the mitochondrial genome (Figure 1D and Table S9). The CNV of Y’-and Ty elements in different strains substantially contributed to the nuclear genome size differences (Figure 1C and Table S8). For example, we observed strain-specific enrichment of full-length Ty1 in the reference *S. cerevisiae* S288c, Ty4 in the South American *S. paradoxus* UFRJ50876 and Ty5 in the European *S. paradoxus* CBS432 whereas no full-length Ty was found in the Malaysian *S. cerevisiae* UWOPS03-461.4 (Table S6). Similarly, >30 copies of Y’-element were found in the *S. cerevisiae* SK1 but none in the Far East Asian *S. paradoxus* N44 (Table S5). As for the size variation among mitochondrial genomes, the dynamic distribution of group I and group II introns (in *COB1*, *COX1* and *rnl*) have clearly played an important role (Figure D and Table S9-S10). Despite the large-scale interchromosomal rearrangements in one *S. cerevisiae* (UWOPS03-461.4) and two *S. paradoxus* (UFRJ50816 and UWOPS91-917.1) strains, all the 12 strains maintained 16 nuclear chromosomes with one centromere on each chromosome, in contrast to the chromosome reduction observed in other post-whole genome duplication (post-WGD) *Saccharomycotina* yeasts (Gordon et al. 2011).

### Contrasting molecular evolutionary rates and diversification timescales between *S. cerevisiae* and *S. paradoxus*

A robust phylogeny is a prerequisite for making reliable evolutionary inferences. We used a concatenated multi-loci matrix of 4,531 one-to-one orthologous nuclear genes to construct a maximum likelihood (ML) phylogenetic relationship of the 12 strains, together with six other *Saccharomyces sensu stricto* species as outgroups. The resulting phylogeny is consistent with our prior knowledge about these strains (Figure 1E). In summary, the seven *S. cerevisiae* strains and the five *S. paradoxus* strains were unambiguously clustered into their respective species clades, which together formed a monophyletic group separated from the outgroup species. The *S. paradoxus* clade was further partitioned into two early-diversified continental groups: the Eurasian group represented by CBS432 (European) and N44 (Far East Asian), and the American group represented by UWOPS91-917.1 (Hawaiian), YPS138 (North American) and UFRJ50816 (South American). The phylogenetic relationship presented here is highly robust, as all the internal nodes have 100% fast-bootstrapping support. We also constructed individual ML gene trees for all the 4,531 one-to-one orthologs and summarized them into a single coalescent-based consensus species tree. This revealed exactly identical topology as the concatenated tree with a normalized quartet score of 0.92. Taken together, these results suggest the inferred phylogeny is highly robust, thereby laying a solid foundation for our downstream evolutionary analysis.

In addition to the topology, we also examined the branch lengths of this phylogenetic tree as well as the chronogram generated by molecular dating to gain insights into the evolutionary rates and timescales of the two species. We found the entire *S. cerevisiae* lineage to have evolved faster than the *S. paradoxus* lineage as indicated by their overall longer branch lengths (from the common ancestor of the two species to each tip of the tree) (Figure 1E). We further confirmed such rate difference in molecular evolution by Tajima’s relative rate test (Tajima 1993) for all *S. cerevisiae* versus *S. paradoxus* strain pairs by using *S. mikatae* as the outgroup (*p*-value < 1E-5 for all pairwise comparisons). The molecular dating analysis (Figure S2) suggests the *S. cerevisiae* strains have diversified much more recently than their *S. paradoxus* counterparts, which was further supported by our synonymous substitution rate (dS) calculation (Figure S3). The cumulative diversification time for the five *S. paradoxus* strains is 3.88 times of that for the seven *S. cerevisiae* strains, suggesting a much longer time span for accumulating evolutionary changes in *S. paradoxus* during its diversification.

### Explicit partitioning of nuclear chromosomes into cores, subtelomeres and chromosome-ends

Conceptually, the linear nuclear chromosomes can be partitioned into three distinct domains: internal chromosomal cores, interstitial subtelomeres and terminal chromosome-ends. However, the boundaries between these domains had never been explicitly demarcated due to the lack of a rigid subtelomere definition. Here, we capitalize on the cross-comparison of multiple complete genome assemblies to strictly define yeast subtelomeres for the two closely related species that we studied. For each subtelomere, we located its proximal boundary based on the sudden loss of synteny conservation on the corresponding chromosome across the 12 strains and demarcated its distal boundary using the yeast-specific telomere-associated sequences, i.e. the core X-and Y'-elements (See Materials and Methods for details) (Figure S4). The partitioning for the left arm of chromosome I (chrI-L) is illustrated in Figure 2. Note that the strict gene synteny conservation is immediately lost after the *GDH3* gene, which marks the boundary between the chromosomal core and the subtelomere for this chromosome arm (Figure 2). All chromosomal cores, subtelomeres, and 358 out of 384 of chromosome-ends across the 12 strains were thus defined (Table S11-13 and Supplementary data 1-2). For the remaining 26 chromosome-ends, neither core X/Y'-elements nor telomeric repeats (TG_1-3_) could be found, suggesting incomplete assemblies.

**Figure 2.**
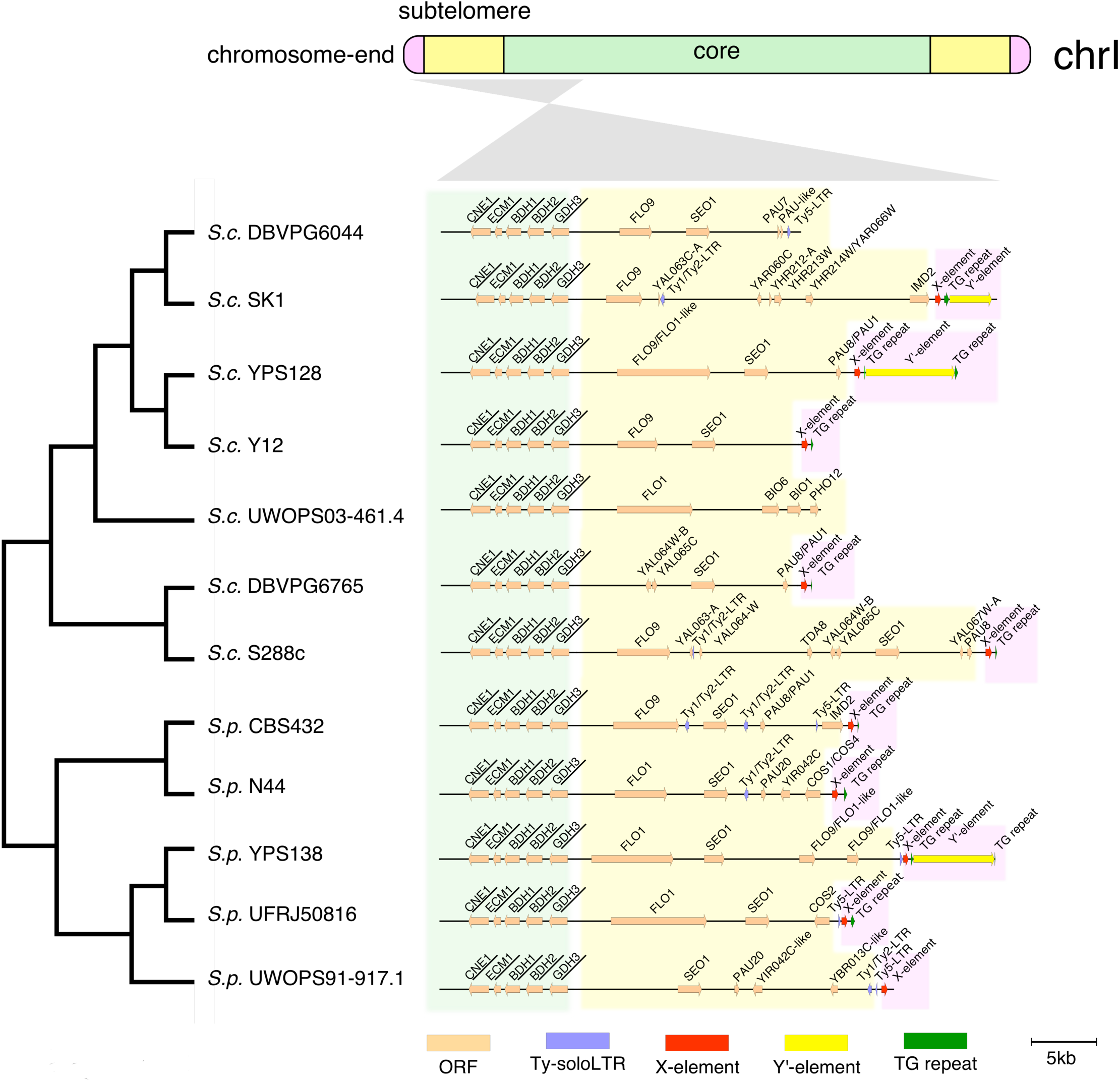
Explicit chromosome partitioning for nuclear chromosomes. In this illustrated example, we partitioned the left arm of chromosome 1 (chrI-L) into the core (light green), subtelomere (light yellow) and chromosome-end (pink) based on synteny conservation and yeast telomere associated sequences (the core X-and Y’-elements). The cladogram (left side) depicts the phylogenetic relationship of the 12 strains, while gene maps (right side) illustrate syntenic conservation in the core region with gene names within syntenic block underlined. Genes are colored in light orange, Ty-related features (all soloLTRs in the plotted region) in light purple, core X-elements, Y’-elements and TG_1-3_-repeats are shown in red, yellow and green respectively.

As a validation of our chromosome partitioning, all yeast genes defined as essential in the *S. cerevisiae* strain S288c fell into the chromosomal cores in all strains and all known subtelomeric duplication blocks in S288c (<u>http://www2.le.ac.uk/colleges/medbiopsych/research/gact/images/clusters-fixed-large.jpg</u>) were fully enclosed in our defined S288c subtelomeres. Furthermore, the genes from our defined subtelomeres show consistently higher rates of molecular evolution and CNV accumulation than those from the cores (Wilcoxon rank sum test, *p*-value = 3.66E-3 for the dN/dS comparison within *S. paradoxus* and *p*-value <2.2E-16 for all the other comparisons) (Figure S5). All these observations fit well with the known biological properties of the cores and subtelomeres. Admittedly, we may underestimate the size of subtelomeres for those chromosomes with extremely high synteny conservation extending all the way to the chromosome-ends (exemplified in the later section). However, we argue that such terminal regions with high synteny conservation deviate from the highly dynamic nature of typical subtelomeres and should therefore be classified as cores.

We assigned the orthologous relationships between our defined subtelomeres from different strains based on the ancestral chromosomal identity of the chromosomal cores that they are attached to, which accounts for the large-scale interchromosomal rearrangements that have occurred in some strains (Table S12). For example, we named the subtelomere located on the right arm of chrXI in UWOPS03-461.4 as the “chr07-L subtelomere” since this subtelomere, together with its flanking chromosomal core, came from the left arm of the ancestral chromosome 7 (chr07-L) (Figure S6). Such accurately assigned subtelomere orthology together with our explicit chromosome partitioning allows us to treat each chromosomal domain separately and to have in-depth examination of their respective evolutionary dynamics.

### Contrasting patterns of structural rearrangements between the two species in chromosomal cores

Structural rearrangements can be balanced (e.g. inversion, reciprocal translocation, and transposition) or unbalanced (e.g. large novel insertion, deletion, and duplication) depending on whether the actual amount of genetic material is affected (Feuk et al. 2006). We systematically identified both types of structural rearrangements in the chromosomal cores of the 12 strains, with a focus on events in which protein-coding genes are involved. The identified structural rearrangements were further mapped back to the strain phylogeny to reconstruct their evolutionary history.

We identified 35 balanced rearrangements in total, including 26 inversions, six reciprocal translocations, two transpositions and one massive rearrangement (Figure 3A and Supplementary data 3). All but one event occurred after the onset of diversification in each species, with most events occurring in the *S. paradoxus* lineage and very few in the *S. cerevisiae* lineage. Even when factoring in the difference in the cumulative diversification time of these two lineages, *S. paradoxus* still shows faster accumulation of balanced rearrangements than *S. cerevisiae*, with a 1.93-fold difference in rates. Of the 26 inversions, six are tightly packed into a ~200 kb region on chrVII of the South American *S. paradoxus* UFRJ50816, indicating a strain-specific inversion hotspot (Figure 3B). Another three inversions occurring independently in two *S. cerevisiae* (SK1 and YPS128) and one *S. paradoxus* (UFRJ50816) strains are located in a known “flip-flop” region surrounded by two inverted homologous segments (Philippsen et al. 1997; Wei et al. 2007; KH Wolfe, personal communication), suggesting a potential scenario of balancing selection acting on this inversion (Supplementary data 3). All the six reciprocal translocations occurred in *S. paradoxus* strains, with five in UFRJ50816 (South American) and one in UWOPS91-917.1 (Hawaiian) (Figure 3C). One transposition is shared by all the seven *S. cerevisiae* strains while the other likely occurred in the common ancestor of the North American (YPS138) and South American (UFRJ50816) *S. paradoxus*. The interchromosomal rearrangement found in the Malaysian *S. cerevisiae* UWOPS03-461.4 is particularly striking, in which chrVII, chrVIII, chrX, chrXI, and chrXIII were completely reshuffled, confirming a recent observation for this strain based on chromosomal contact data (Marie-Nelly et al. 2014) (Figure 3C). We coined the term “massive rearrangement” to describe such dramatic genome reconfiguration, as it cannot be explained by typical reciprocal translocations. This may result from a single catastrophic event resembling the chromothripsis observed in tumor cells (Stephens et al. 2011; Zhang et al. 2013) or from multiple independent events separated in time. The massive rearrangement in the Malaysian *S. cerevisiae* UWOPS03-461.4 and the rapid accumulation of inversions and translocations in the South American *S. paradoxus* UFRJ50816 resulted in extensively altered genome configurations, which explain the reproductive isolation of these two lineages (Liti et al. 2006; Cubillos et al. 2011). Finally, as previously observed in yeasts with larger divergence scales (Fischer et al. 2000; Kellis et al. 2003), the breakpoints of these balanced rearrangements are clearly associated with tRNAs and Tys, highlighting the roles of these elements in triggering genome instability.

**Figure 3.**
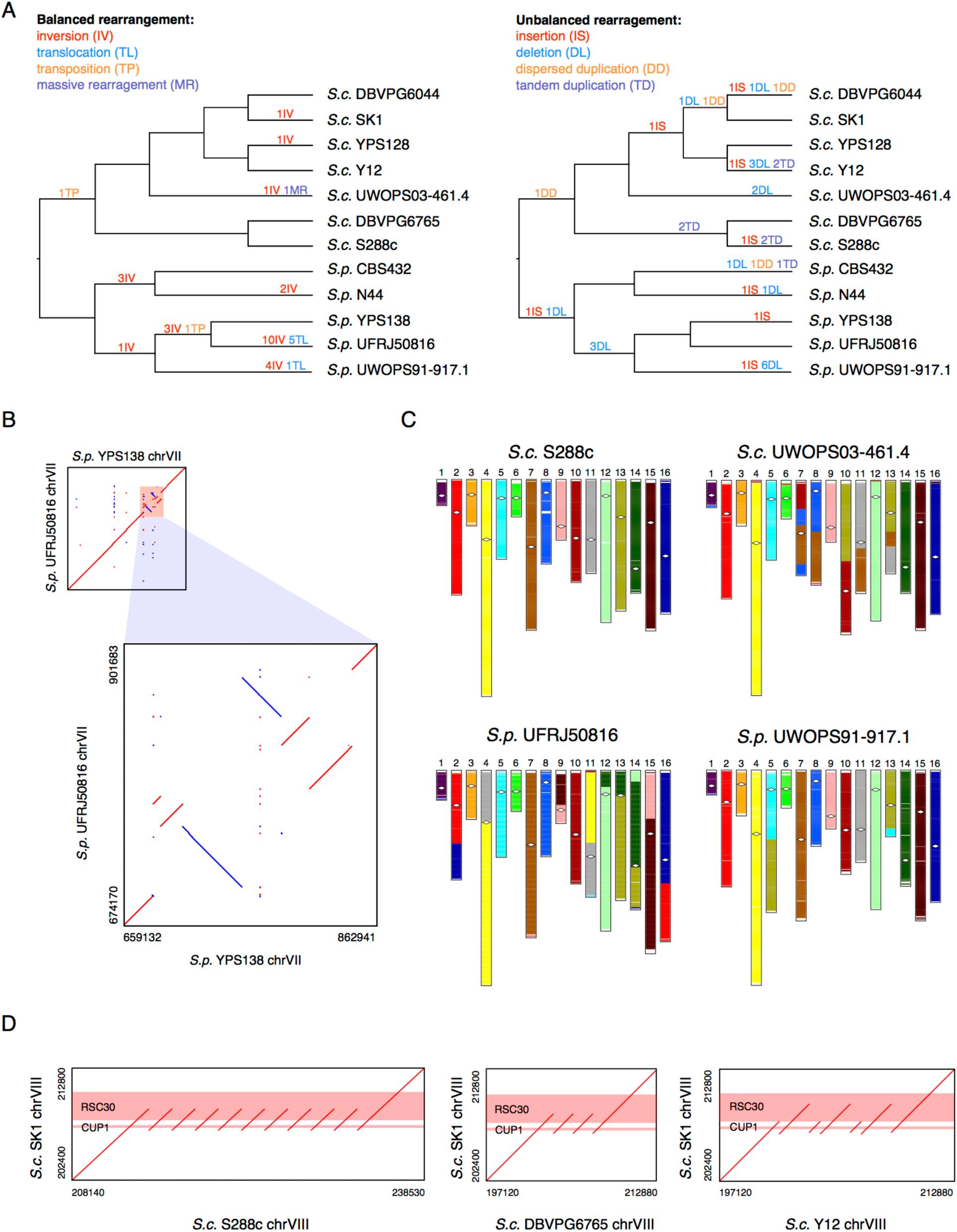
Structural rearrangements in the nuclear chromosome cores. (A) Balanced (left side) and unbalanced (right side) structural rearrangements occurred along the evolutionary history of the 12 strains. (B) The six clustered inversions on chrVII of the South American *S. paradoxus* UFRJ50816. (C) Genome organization of the strains UWOPS03-461.4, UFRJ50816 and UWOPS91-917.1 relative to that of S288c. The strain S288c is free from large interchromosomal rearrangement, and could therefore represent the ancestral genome organization. White diamonds indicate centromere position. (D) Dotplots showing tandem duplications of the *CUP1*-*RSC30* locus in S288c, DBVPG6765, and Y12. The genic regions of *CUP1* and *RSC30* gene are highlighted in red.

Considering unbalanced structural rearrangements, we identified eight novel insertions, 19 deletions, four dispersed duplications and at least seven tandem duplications (Figure 3A and Supplementary data 4). There are another two cases (one dispersed duplication and one insertion/deletion) of which the evolutionary history cannot be confidently determined due to potentially multiple independent origins or secondary deletions (Supplementary data 4). Although these numbers can be slightly underestimated given that we only count unambiguous cases in our analysis, our identified unbalanced structural rearrangements clearly outnumbered those balanced ones, as recently found in *Lachancea* yeasts (Vakirlis et al. 2016). In contrast to the balanced rearrangements, we found the unbalanced ones to be much more evenly distributed across the strains in both species despite the much shorter cumulative diversification time of *S. cerevisiae*. We estimated that the accumulation rate of unbalanced rearrangements in *S. cerevisiae* is 4.6 times of that in *S. paradoxus* during the respective diversification of these two species. The unbalanced structural rearrangements also occurred at a much smaller genomic scale compared to the balanced ones, with at most a few genes involved in each event. We found that the breakpoints of these unbalanced rearrangements (except for the tandem duplications) were also associated with Tys and tRNAs, echoing our observation for balanced rearrangements. The genes *CUP1*, *RSC30*, *ENA1/2/5*, and *TDH3* are involved in tandem duplications, with the case at the *CUP1*-*RSC30* locus being especially intriguing. The *CUP1* and *RSC30* genes were tandemly duplicated in three *S. cerevisiae* strains (DBVPG6765, S288c and Y12) with different duplication segments, at least between Y12 and the other two strains (Figure 3D). This likely suggests a scenario of convergent evolution driven by selection for copper tolerance in independently domesticated beverage producing lineages as previously suggested (Warringer et al. 2011). Finally, we found genes involved in unbalanced rearrangements to be significantly enriched for gene ontology (GO) terms related to the binding, transporting and detoxification of metal ions (e.g. Na^+^, K^+^, Cd^2+^ and Cu^2+^) (Table S14), suggesting these events likely to be adaptive.

### High resolution view of structural plasticity and evolutionary dynamics in subtelomeres

Our complete assemblies and explicitly defined subtelomere boundaries allowed us to examine the structural plasticity and evolutionary dynamics of subtelomeres with unprecedented resolution. The subtelomere sizes are highly variable across different strains and chromosome arms, ranging from 0.13 to 76 kb (median = 15.6 kb) (Figure 4A and Supplementary data 2). While the very short subtelomeres (e.g. the chr04-R and chr11-L subtelomeres) can be explained by the widespread high degree of synteny conservation extending all the way to the chromosome-ends, those exceptionally long subtelomeres can instead be caused by multiple mechanisms. For example, the chr15-R subtelomere of the European/Wine *S. cerevisiae* DBVPG6765 has been drastically elongated by a 65 kb genomic segment that was horizontally transferred from *Torulaspora microellipsoides* (Marsit et al. 2015) (Figure 4B and Figure S7A). The chr07-R subtelomere of the European *S. paradoxus* CBS432 was extended by the tandem duplications of *MAL31*-like and *MAL33*-like genes as well as the addition of a terminal segment containing the *ARR* cluster (Figure 4C and Figure S7B). The chr15-L subtelomere of the South American *S. paradoxus* UFRJ50816 increased its size by duplications of subtelomeric segments from two other chromosomes (Figure 4D and Figure S7C). Inversions have also occurred in subtelomeres, including one affecting the *HMRA1*-*HMRA2* gene cluster in UFRJ50816 (Figure 4E) and another affecting an *MAL11*-like gene in the European *S. paradoxus* CBS432 (Figure 4F).

**Figure 4.**
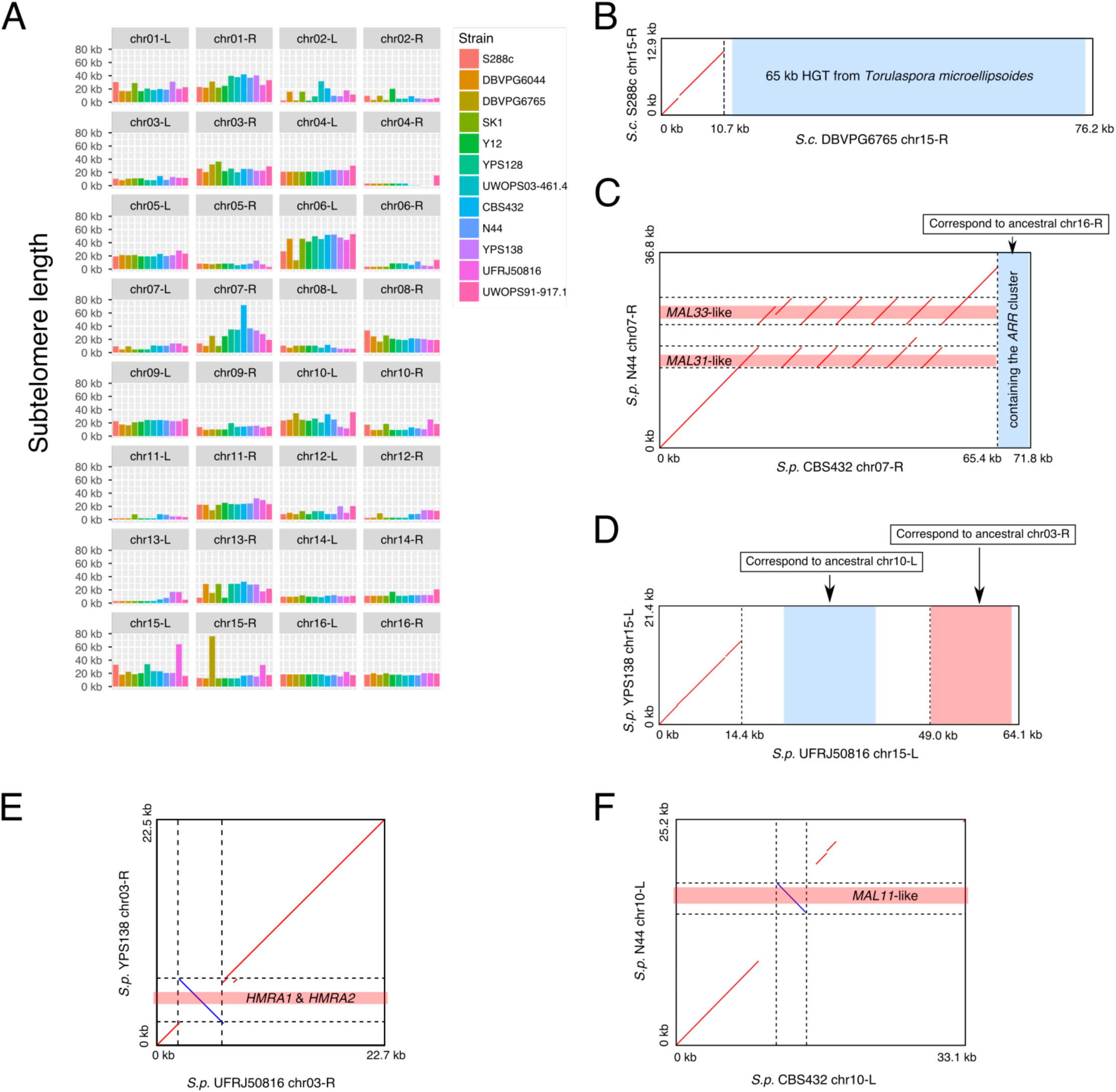
Subtelomere length plasticity and structural rearrangements. (A) Length variation of the 32 orthologous subtelomeres across the 12 strains. The orthologous subtelomeres are assigned based on the ancestral chromosomal identity of the core regions that they are attached to. (B) Dotplot for the chr15-R subtelomere comparison between the *S. cerevisiae* DBVPG6765 and S288c. The much longer DBVPG6765 chr15-R subtelomere is explained by a previously reported 65-kb eukaryote-to-eukaryote horizontal gene transfer (HGT) event. (C) Dotplot for the chr07-R subtelomere comparison between the *S. paradoxus* CBS432 and N44. The much longer chr07-R subtelomere in CBS432 is explained by a series of tandem duplications of the *MAL31*-like and *MAL33*-like genes and an addition of the *ARR*-containing segment from the ancestral chr16-R subtelomere. (D) Dotplot for the chr15-L subtelomere comparison between the *S. paradoxus* UFRJ50816 and YPS138. The much longer chr15-L subtelomere in UFRJ50816 is explained by the relocated subtelomeric segments from the ancestral chr10-L and chr03-R subtelomeres respectively. (E) Dotplot for the chr03-R subtelomere comparison between the *S. paradoxus* UFRJ50816 and YPS138 reveals an inversion occurred at the *HMR* locus in UFRJ50816. (F) Dotplot for the chr03-R subtelomere comparison between the *S. paradoxus* CBS432 and N44 reveals an inversion occurred at an *MAL11*-like locus in CBS432. Please note that the region coordinates for B-F are based on the extracted subtelomeric regions rather than the full chromosomes.

The enrichment of segmental duplications via ectopic sequence reshuffling is a common feature of eukaryotic subtelomeres. Here, we identified such subtelomeric duplication blocks based on pairwise comparisons of different subtelomeres within the same strain (Supplementary data 5). Figure 5A illustrates an example showing duplication blocks shared among three subtelomeres (from chr01-L, chr01-R and chr08-R) in the *S. cerevisiae* reference strain S288c. In total, we identified 173 pairs of subtelomeric duplication blocks across the 12 strains, with 8-26 pairs for each strain (Table S15). Among the 16 pairs of subtelomeric duplication blocks previously identified in S288c, we recaptured all the 12 major pairs, leaving the remaining four pairs too small to pass our filtering criteria. Interestingly, the Hawaiian *S. paradoxus* UWOPS91-917.1 has the most subtelomeric duplication blocks and half of them are strain-specific, suggesting unique subtelomeric evolution in this strain. For all duplication blocks, we noticed that the duplicated segments always maintained the same centromere-telomere orientation, supporting a mechanism of double-strand break (DSB) repair as previously suggested in other species (Linardopoulou et al. 2005; Fairhead and Dujon 2006). We further summarized those 173 pairs of duplication blocks based on the orthologous subtelomere pairs that were involved. This led to 75 unique duplicated subtelomere pairs, 59 of which are new compared to what was previously identified in S288c. We found 31 (41.3%) of these unique pairs to be shared between strains or even between species with highly dynamic strain-sharing patterns, most (87.1%) of which cannot be explained by the strain phylogeny (Figure 5B and Supplementary data 6). This suggests a constant gain and loss process of subtelomeric duplication throughout evolutionary history.

**Figure 5.**
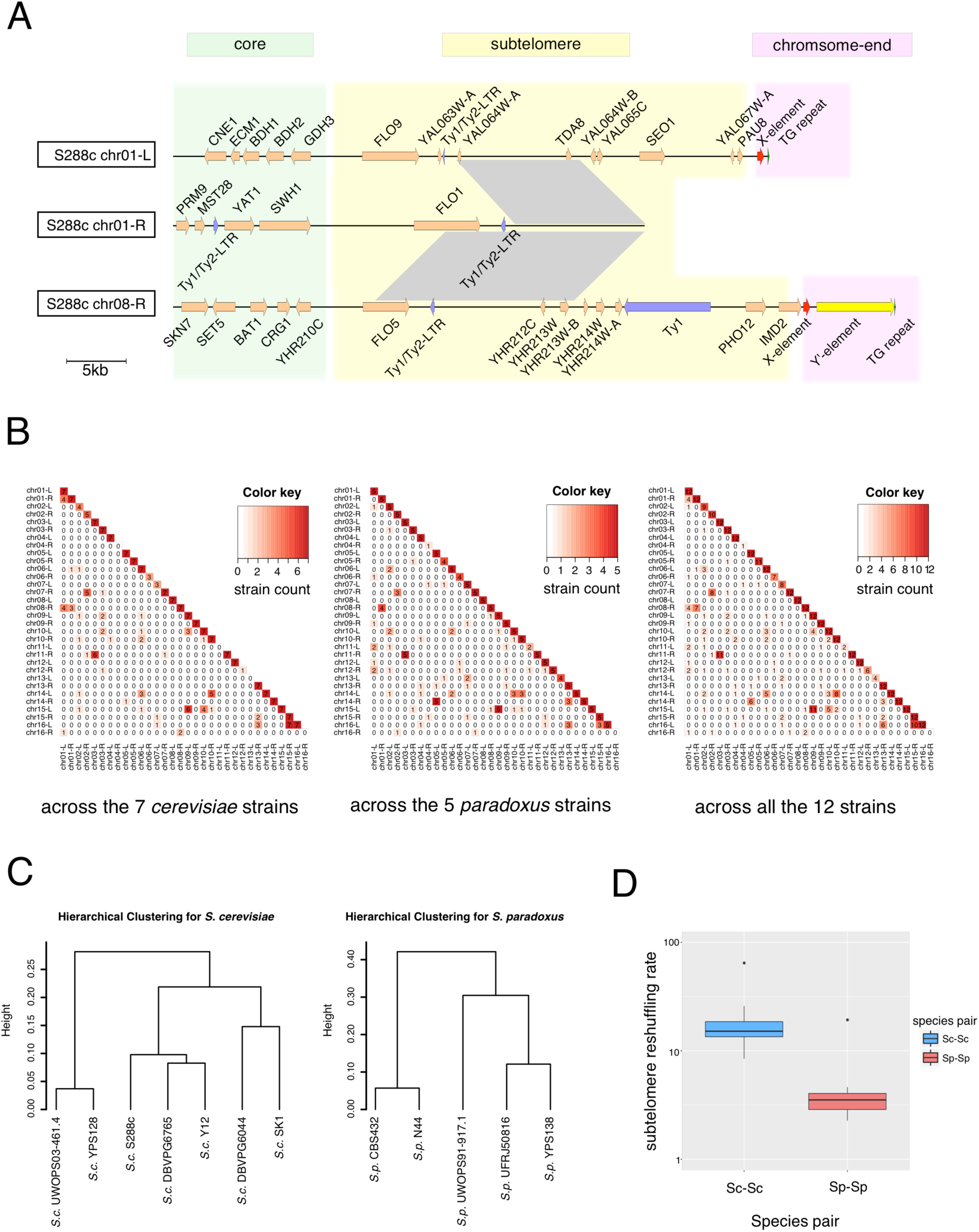
Evolutionary dynamics of subtelomeric duplications. (A) An example of subtelomeric duplication blocks shared among the chr01-L, chr01-R and chr08-R subtelomeres in *S. cerevisiae* S288c. The grey blocks denote their shared homologous regions with >= 90% nucleotide sequence identity. (B) Subtelomeric homology signal shared across the seven *S. cerevisiae* strains, the five *S. paradoxus* strains and all the 12 strains (from left to right respectively). For each pairwise subtelomere combination, the number of strains showing strong sequence homology (BLAT score >= 5000 and identity >= 90%) for this specific subtelomere-pair was counted and visualized using heatmaps. (C) Hierarchical clustering based on proportion of conserved orthologous subtelomeres in cross-strain comparisons within *S. cerevisiae* and within *S. paradoxus* respectively. (D) Subtelomere reshuffling rates within *S. cerevisiae* (Sc-Sc) and within *S. paradoxus* (Sp-Sp). The y-axis is in a log-10 scale.

Given such highly dynamic nature of subtelomeric reshuffling, we investigated to what extent those orthologous subtelomeres could reflect the intra-species phylogeny. We measured the proportion of conserved orthologous subtelomeres in all strain pairs within the same species and performed hierarchical clustering accordingly (Figure 5C). While the clustering on *S. paradoxus* strains correctly recapitulated their true phylogeny, our parallel analysis in *S. cerevisiae* turned out to be very noisy, with only the relationship of the most recently diversified strain pair (DBVPG6044 vs. SK1) being correctly recovered. The fact that the distantly related European/Wine (DBVPG6765) and Sake (Y12) *S. cerevisiae* strains were clustered together indicates likely convergent subtelomere evolution during their respective domestication. The proportion of conserved orthologous subtelomeres between *S. cerevisiae* strains (56.3%-81.3%) is comparable to that between *S. paradoxus* strains (50.0%-81.3%), despite the much smaller diversification timescales of *S. cerevisiae*. Therefore, the contrasting result between the two species from our clustering analysis implies potentially more rapid subtelomeric reshuffling in *S. cerevisiae* than in *S. paradoxus* during their respective diversifications. Indeed, we found a 4.3-fold rate difference in subtelomere reshuffling between the two species (Wilcoxon rank sum test, *p*-value = 2.52E-4; see Methods for details about rate estimation) (Figure 5D), which explains the substantial erosion of true phylogenetic signals in the subtelomere evolution of *S. cerevisiae*. Interestingly, the most recently diversified strain pairs in both species (DBVPG6044 vs. SK1 in *S. cerevisiae* and YPS138 vs. UFRJ50816 in *S. paradoxus*) showed exceptionally high subtelomeric reshuffling rates compared with the other strain pairs within the same species (the outliers in Figure 5D), suggesting possible accelerated subtelomere reshuffling in the incipient phase of strain diversification. The frequent reshuffling of subtelomeric sequences can have drastic impacts on subtelomeric gene content both qualitatively and quantitatively. For example, four genes (*PAU3*, *ADH7*, *RDS1*, and *AAD3*) were lost in the Sake *S. cerevisiae* (Y12) due to a single chr08-L to chr03-R subtelomeric duplication event (Figure S8). Therefore, the more rapid subtelomere reshuffling in *S. cerevisiae* could have important functional implications.

### Non-canonical chromosome-end structures at native telomeres

*S. cerevisiae* chromosome-ends are characterized by two telomere associated sequences: the core X-and Y'-elements (Louis 1995). The core X-element is present in almost all chromosome-ends, whereas the Y'-element is highly variable in terms of both presence/absence and copy numbers across different chromosome-ends and strains. The typical structure of *S. cerevisiae* chromosome-ends can be summarized into two general types: 1) with a single core X-element but no Y’-elements and 2) with a single core X-element followed by one or several distal Y'-elements (Louis 1995). *S. paradoxus* chromosome-ends also contain core X-and Y’-elements (Liti et al. 2009b), but their detailed structures have not been systematically characterized in a genome-wide fashion due to the lack of complete assemblies. Across our 12 strains, most (~85%) chromosome-ends have one of the two typical structures previously characterized in *S. cerevisiae* but we also discovered several non-canonical structures that have not been described before (Table S13). For example, we found several examples of tandem duplications of the core X-element in both species. Such core X-element duplications are unlikely to be assembly artifacts given that we also detected them in the *S. cerevisiae* reference genome (chrVIII-L and chrXVI-R) with degenerated proximal copies. In most cases, the proximal duplicated copies of the core-X element were degenerated but we also found two examples where intact duplicated copies were retained: the chrXII-R end in the Sake *S. cerevisiae* Y12 and the chrIII-L end in the European *S. paradoxus* CBS432. The latter case is especially striking, where six copies (including three complete ones) of the core X-element were tandemly arranged. Even more surprisingly, we discovered five chromosome-ends consisting of only Y'-element (one or more copies) but no core X-element, despite the importance of the core X-element in maintaining genome stability (Marvin et al. 2009a, 2009b). For example, the chrV-L ends in *S. cerevisiae* DBVPG6044 and SK1 have one and three Y'-elements respectively without any trace of the core X-elements. The discoveries of these non-canonical chromosome-end structures offer a new paradigm to investigate the functional role of the core X-elements.

### Lineage-specific structural rearrangements and introgression in the *S. paradoxus* mitochondrial genomes

The mitochondrial genome constitutes a natural genetic compartment that is replicated and transmitted independently from the nuclear genome. Despite its pivotal evolutionary and functional importance, sequencing and assembling the yeast mitochondrial genome has always been challenging due to its highly repetitive and AT-rich genome composition. Obtaining complete mitochondrial genome assemblies from our long-read sequencing gave us a great opportunity to investigate the structural dynamics of the mitochondrial genome with high resolution and accuracy. We found a high degree of collinearity in *S. cerevisiae* mitochondrial genomes for all pairwise comparisons, even between the most distantly related strains (e.g. DBVPG6044 vs. S288c) (Figure 6A). In contrast, the *S. paradoxus* mitochondrial genomes show lineages-specific structural rearrangements in several strains. The two Eurasian strains (CBS432 and N44) share a transposition of the entire *COX3*-*rnpB*-*rns* segment, in which *rns* was further inverted either before or after the transposition (Figure 6B-D). Independent tandem duplications of the *OLI1-VAR1* segment and *rnpB* were found in the South American (UFRJ50816) and Hawaiian (UWOPS91-917.1) *S. paradoxus* respectively (Figure 6E and Table S9). In addition, it seems that the *COB* gene was recently transposed to its current mitochondrial genomic position in *S. cerevisiae* and *S. paradoxus* prior to the divergence of the two species given the gene orders in the two outgroups.

**Figure 6.**
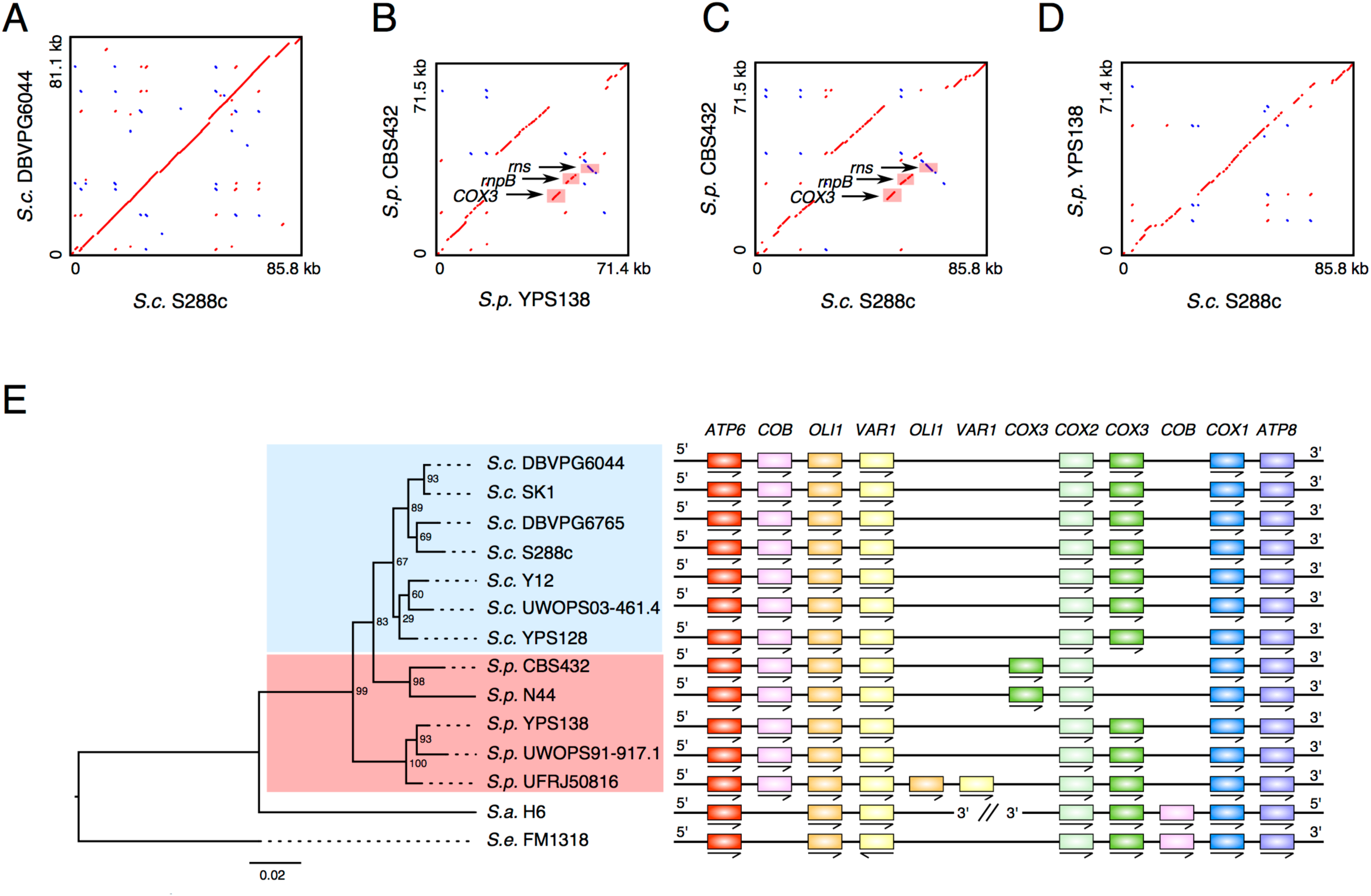
Comparative mitochondrial genomics. (A) Pairwise comparison for the mitochondrial genome of S288c and DBVPG6044 from *S. cerevisiae*. (B) Pairwise comparison for the mitochondrial genome of CBS432 and YPS138 from *S. paradoxus*. The transposition of *COX3* and the inversion of *rns* are highlighted in the plot. (C) Pairwise comparison for the mitochondrial genome of *S. cerevisiae* S288c and *S. paradoxus* CBS432. The transposition of *COX3* and the inversion of *rns* are highlighted in the plot. (D) Pairwise comparison for the mitochondrial genome of *S. cerevisiae* S288c and *S. paradoxus* YPS138. (E) Protein-coding gene arrangement in the mitochondrial genome across the 12 sampled strains. The phylogenetic tree shown on the left is based on the concatenated protein sequence of the six one-to-one mitochondrial genes (*ATP6*, *ATP8*, *COB*, *COX1*, *COX2*, and *COX3*). The numbers at the internal nodes are the rapid bootstrap value showing statistical supports for the corresponding node. The detailed gene arrangement in each strain is shown on the right.

The phylogenetic tree inferred from the six one-to-one mitochondrial orthologs (*ATP6*, *ATP8*, *COB*, *COX1*, *COX2* and *COX3*) deviates from the phylogeny based on nuclear genes, although the relationships between those most closely related strains (e.g. DBVPG6044 vs. SK1 and DBVPG6765 vs. S288c in *S. cerevisiae* as well as CBS432 vs. N44 in *S. paradoxus*) are the same. The low topology consensus across different gene loci (normalized quartet score = 0.594) suggests a highly heterogeneous phylogenetic history of mitochondrial genes across different loci. Together with the drastically dynamic presence/absence pattern of mitochondrial group I and group II introns (Table S10), this supports the idea of extensive cross-strain recombination in yeast mitochondrial evolution (Wu et al. 2015a). According to this mitochondrial phylogeny, the Eurasian *S. paradoxus* lineage (CBS432 and N44) was clustered together with the seven *S. cerevisiae* strains before joining with the other *S. paradoxus* strains, which reinforces the argument for mitochondrial introgression from *S. cerevisiae* to the Eurasian *S. paradoxus* lineage (Wu and Hao 2015) (Figure 6E). In addition, we noticed that the *COX3* gene in the South American *S. paradoxus* UFRJ50816 started with GTG rather than the typical ATG start codon in our assembly, which was further confirmed by the Illumina reads. This suggests either an adoption of an alternative nearby ATG start codon (e.g. the one 45 bp downstream) or a rare case of near-cognate start codon as used by bacteria (Blattner et al. 1997; Cole et al. 1998) and *Candida* yeast (Abramczyk et al. 2003).

### Fully resolved structural rearrangements illuminate complex phenotypic traits

Structural rearrangements are expected to account for a substantial fraction of phenotypic variation but the lack of complete assemblies have prevented a deep understanding of structural variation–phenotype associations. Here we used the *CUP1* locus and *ARR* cluster as case studies to illustrate how the fully resolved structural rearrangements based on complete assemblies can illuminate complex phenotypic traits. The *CUP1* gene encodes a copper scavenging short metallothionein that keeps the intracellular level of free copper extremely low and mediates copper tolerance. Across our 12 strains, this gene was tandemly amplified into four, seven, and 11 copies in three *S. cerevisiae* strains (DBVPG6765, Y12 and S288c respectively) while maintaining the ancestral single copy configuration in all the other strains (Figure 7A). Consistent with previous observations (Warringer et al. 2011), such copy number variation matched well with the strain growth rates in high copper concentration conditions (CuCl2: 0.38 mM) (Figure 7B). In general, higher copy number of *CUP1* translates into faster growth (i.e. shorter generation time) in copper, although Y12 with seven copies appears to grow slightly faster than S288c with 11 copies (including one pseudogene copy), which could be explained by differences in the amplification segment and/or the overall genetic backgrounds. The *ARR* cluster contains three consecutive subtelomeric genes (*ARR1*, *ARR2* and *ARR3*) that function collectively to provide arsenic resistance. Despite their tricky genomic locations (only a few kb from the core-X element), we successfully characterized the exact genomic arrangement of the *ARR* cluster in all the 12 strains (Figure 7C). Consistent with our previous estimates based on read mapping coverage (Bergström et al. 2014), the *ARR* cluster was duplicated in the European *S. paradoxus* CBS432 while completely lost in two *S. cerevisiae* (SK1 and UWOPS03-461.4) and two *S. paradoxus* (N44 and UWOPS91-917.1) strains (Figure 7C). Our growth rate assay confirmed the link between *ARR* cluster loss and extreme susceptibility to arsenic (3 mM arsenite, As[III]) (Figure 7D). Despite having two copies of the *ARR* cluster, CBS432 grew poorly in arsenic. Since no strongly deleterious mutation was detected in either gene or copy, the arsenic sensitivity of CBS432 should derive from its genetic background. The As[III] sensitivity of the South American *S. paradoxus* UFRJ50816 could potentially be explained by the pseudogenization of its *ARR2*, although it is only known to protect against pentavalent arsenic, As[V] (Mukhopadhyay et al. 2000). The genes immediately proximal to the *ARR* cluster are located at the chr16-R subtelomere in all the 12 strains, implying that this subtelomere should be the ancestral location for the *ARR* cluster. In the West African *S. cerevisiae* DBVPG6044, *ARR* became relocated to the chr03-R subtelomere. In the European *S. paradoxus* CBS432, the two duplicated *ARR* clusters were redistributed to the chr02-R and chr07-R subtelomeres respectively. In the two American *S. paradoxus* (YPS138 and UFRJ50816), the *ARR* cluster was shifted to the ch13-L subtelomere with an inverted orientation.

**Figure 7.**
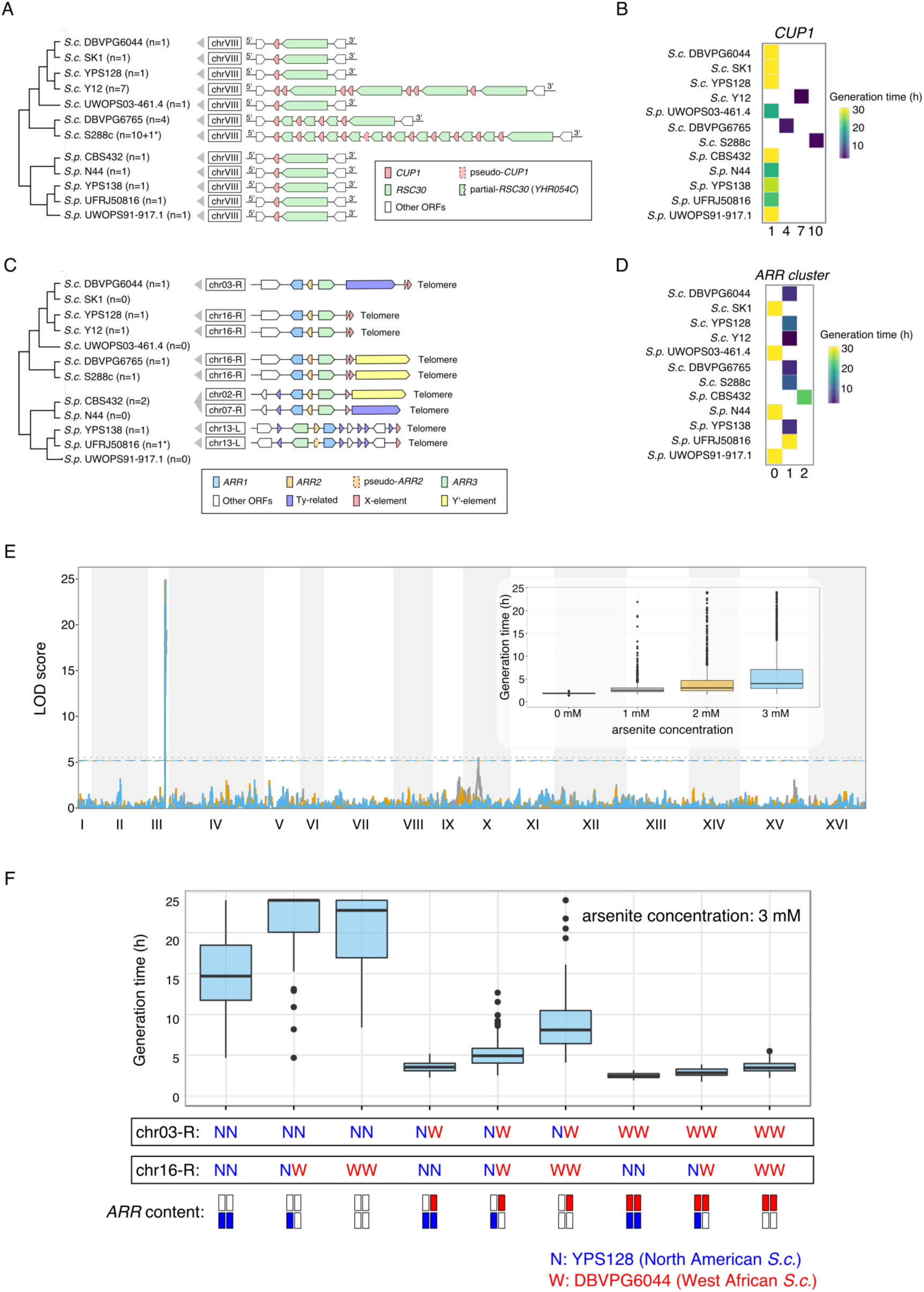
Structural rearrangements account for complex phenotypic variation. (A) Copy number and gene arrangement of the *CUP* locus across the sequenced strains. The asterisk on the copy number denotes the involvement of pseudogene. (B) Generation time of the 12 strains in high copper concentration condition correlates with *CUP1* copy number. (C) Copy number and gene arrangement of the *ARR* cluster. The asterisk on the copy number denotes the involvement of pseudogene. The subtelomere location of the *ARR* cluster is highly variable. (D) Generation time of the 12 strains in arsenic condition. (E) The rearrangement that relocates the *ARR* cluster to the chr03-R subtelomere in the West African *S. cerevisiae* DBVPG6044 is consistent with the QTL mapping by linkage analysis. This analysis was performed using the recently described phased outbred lines (POLs) technology (Hallin et al. 2016) derived from the North American (YPS128) and West African (DBVPG6044) *S. cerevisiae*. (F) Phenotypic distribution of 826 POLs for generation time in arsenic condition partitioned for genotype positions at the chr03-R and chr16-R subtelomere and inferred copies of *ARR* clusters (underneath the plot).

To quantify how different genomic arrangements of the *ARR* cluster can affect fitness in arsenic (As[III]), we performed quantitative trait locus (QTL) mapping using 826 phased outbred lines (POLs) derived from an advanced intercross of the North American (YPS128) and West African (DBVPG6044) *S. cerevisiae* strains (see Methods for details). The linkage analysis accurately mapped a large-effect QTL at the chr03-R subtelomere (the location of *ARR* in DBVPG6044), but no contribution to arsenic variation from the YPS128 *ARR* on the chr16-R subtelomere (Figure 7E). This profile is consistent with the relocation of an active *ARR* cluster to the chr03-R subtelomere in DBVPG6044 and the presence of deleterious mutations predicted to inactivate the *ARR* cluster in YPS128 (Cubillos et al. 2011; Bergström et al. 2014). The combined effect of genotype and copy number can be fully decomposed by knowing the correct subtelomere structure and segregation pattern (Figure 7F).

## Discussion

The landscape of genetic variation is shaped by multiple evolutionary processes, including mutation, drift, recombination, gene flow, natural selection and demographic history. The combined effect of these different factors can vary considerably both across the genome and between species, resulting in different patterns of evolutionary dynamics. The complete genome assemblies that we generated for multiple representative strains from both domesticated and wild yeasts provide a valuable dataset to explore such patterns with unprecedented resolution.

Considering the dynamics across the genome, it has been observed in many organisms (e.g. fruitfly (Anderson et al. 2008), *Arabidopsis* (Kuo et al. 2006) and human (Linardopoulou et al. 2005)) that eukaryotic subtelomeres usually exhibit exceptional variability in comparison with the internal chromosomal cores. The high evolutionary dynamics of subtelomeres are often manifested by rapid molecular evolution, extensive copy number variation (CNV), and rampant interchromosomal reshuffling, as previously showed in yeasts (Brown et al. 2010; Bergström et al. 2014; Louis and Haber 1990; Fairhead and Dujon 2006; Anderson et al. 2015). Our whole genome comparison both within and between species corroborated all these previous findings and further highlighted the pronounced distinction between the cores and subtelomeres with a focus on structural genome evolution. In contrast to the evolutionarily static chromosomal cores with limited and mostly tractable rearrangement events, subtelomeres showed extreme plasticity and a constant gain and loss of interchromosomal reshuffling, of which the detailed evolutionary history cannot be confidently reconstructed. Such ectopic reshuffling among different subtelomeres can substantially change the content and diversity of the gene repertoire in this highly variable region and even create novel recombinant genes with adaptive potentials (Anderson et al. 2015). Given that the subtelomeric genes are highly enriched in mediating interactions with external environments (e.g. stress response, nutrient uptake and catabolism, and metal/toxin transport) (Ames et al. 2010; Brown et al. 2010; Bergström et al. 2014), it is tempting to speculate that the accelerated subtelomeric evolution at both gene and structural level is at least partially a reflection of selection for evolvability and the capacity for fast adaptation to ecological changes.

While the evolutionary dynamics across the genome is more related to the intrinsic properties of different genomic domains (e.g. cores vs. subtelomeres), external factors such as selection and demographic history hold the key roles in shaping lineage-specific genome dynamics. The ecological niches and recent evolutionary history of *S. cerevisiae* have been associated with human activities, with many strains isolated from human-associated environments like breweries, bakeries and even clinical patients (Liti 2015). Consequently, both natural and artificial selection significantly influenced the genome evolution of these strains, resulting in improved alcoholic fermentation (Fay and Benavides 2005) and enhanced pathogenicity (Muller et al. 2011; Strope et al. 2015), which are adaptive for their respective niches. In addition to introducing novel selection schemes, human activities also help to promote migration, and consequently mixture and crossbreeding of *S. cerevisiae* strains from different geographical locations and ecological niches (Hyma and Fay 2013). Consistent with this notion, previous phylogenetic and population structure analyses on *S. cerevisiae* uncovered many mosaic strains with mixed genetic backgrounds (Liti et al. 2009a). In contrast, the wild-living *S. paradoxus* shows well-differentiated lineages with geographically defined population structure (Koufopanou et al. 2006; Liti et al. 2009a) and partial reproductive isolation between strains from different lineages (Sniegowski et al. 2002; Liti et al. 2006). All currently identified *S. paradoxus* strains were isolated from natural habitats with no evident human interference, which provides an ideal control relative to *S. cerevisiae* to examine the influence of human activities in shaping genome evolution. Here, we summarized the major differences in the evolutionary dynamics of these two species during their respective diversification (Figure 8). In nuclear chromosomal cores, *S. cerevisiae* strains show much lower rate of accumulating balanced structural rearrangements compared with *S. paradoxus* strains. This pattern is likely explained by the mixture and crossbreeding between different *S. cerevisiae* subpopulations during their recent association with human activities, which would considerably impede the fixation of balanced structural rearrangements in different subpopulations. In contrast, the geographical isolation of different *S. paradoxus* subpopulations would be favored for the fixation of balanced rearrangements in different subpopulations (Leducq et al. 2016). As for unbalanced rearrangements in chromosomal cores, we observed an opposite pattern, in which the *S. cerevisiae* strains exhibit higher rate of accumulating such changes than their *S. paradoxus* counterparts. A strong association was further found between genes affected by unbalanced rearrangements and the cellular adaptation to metal ion stress, which collectively indicates a significant role of selection in shaping this pattern. Compared with *S. paradoxus*, *S. cerevisiae* strains are evolving under a much wider spectrum of selection regimes and thus will likely favor accumulating more adaptive changes in strains from different ecological niches. Consistent with this notion, the more rapid interchromosomal reshuffling in *S. cerevisiae* than in *S. paradoxus* is probably also a consequence of selection given the functional importance of subtelomeric genes in promoting adaptation. In both core and subtelomeres, we observed consistently higher rates of molecular evolution and CNV accumulation in *S. cerevisiae* strains, which provide further supports to this argument (Figure S5). In addition, we found that the mitochondrial genomes of the *S. cerevisiae* strains maintained high degrees of collinearity, whereas those of the *S. paradoxus* strains showed lineage-specific structural rearrangements and introgression, suggesting distinct mitochondrial evolution between the two species. Taken together, many of these observed differences between *S. cerevisiae* and *S. paradoxus* reveal the influence of human activities on structural genome evolution.

**Figure 8.**
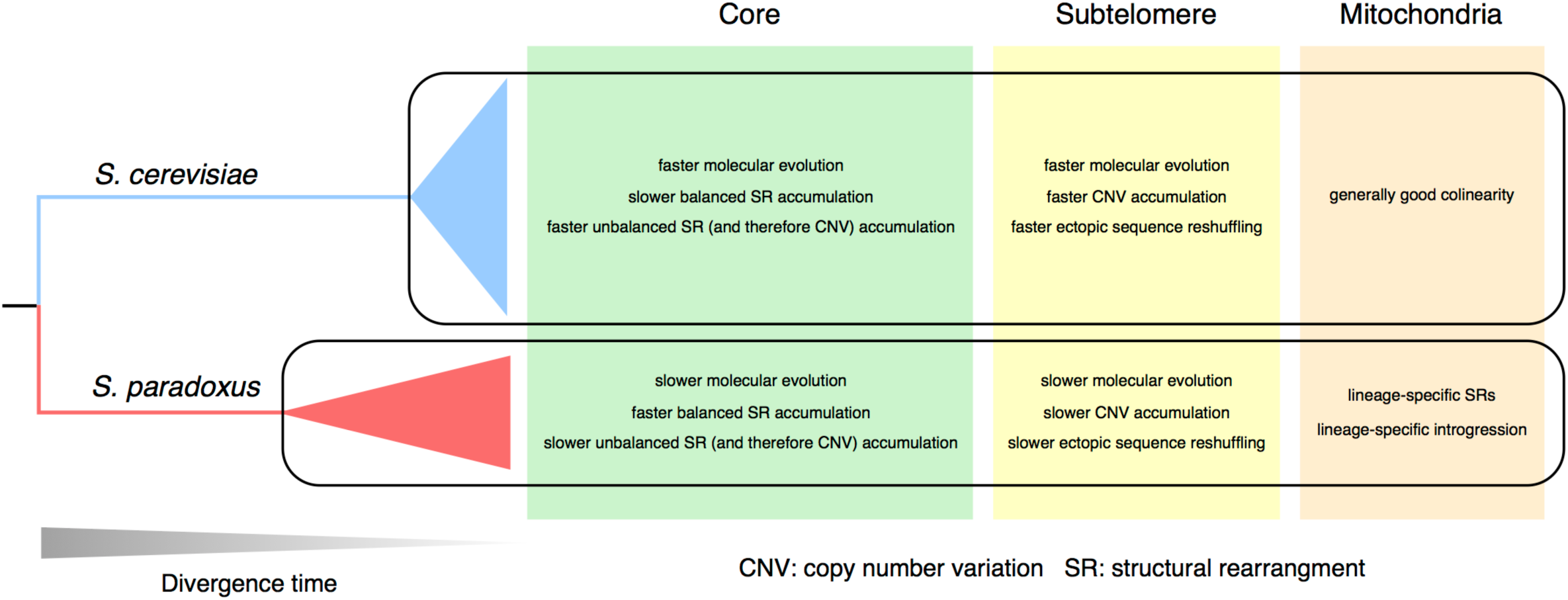
Contrasting evolutionary dynamics across the genome and between species. The evolutionary dynamics of *S. cerevisiae* and *S. paradoxus* during their respective diversifications are summarized with regard to molecular evolution (dN/dS), copy number variation (CNV), balanced and unbalanced structural rearrangements. Note that CNV is the result of unbalanced structural rearrangements.

## Materials and Methods

### Strain sampling, preparation and DNA extraction

Based on previous population genomics surveys (Liti et al. 2009a), we sampled seven *S. cerevisiae* and five *S. paradoxus* strains (all in haploid form or homozygous diploids) to represent the main evolutionary lineages. Our strain sampling also includes the reference strains for *S. cerevisiae* (S288c) and *S. paradoxus* (CBS432) as well as another popular *S. cerevisiae* lab strain (SK1), which were also used for quality control of our sequencing, assembly, annotation and downstream analysis.

All the strains were taken from our strain collection stored at −80°C and cultured on YPD plates. Single colony for each strain was picked and cultured in 5 mL YPD liquid at 30°C 220 rpm overnight. The DNA extraction was carried out using the MasterPure™ Yeast DNA Purification Kit (Epicentre, WI, USA) following the manufacturer's protocol.

### PacBio sequencing and raw assembly

The sequencing center at the Wellcome Trust Sanger Institute (Cambridge, UK) performed library preparation and sequencing using the PacBio Single Molecule, Real-Time (SMRT) DNA sequencing technology. The raw PacBio reads were generated by the PacBio RS II platform with the P6-C4 chemistry and were processed and assembled using the standard SMRT analysis pipeline (v2.3.0). The *de novo* assembly was carried out following the standard hierarchical genome-assembly process (HGAP) assembly protocol with Quiver polishing (Chin et al. 2013).

### Assembly evaluation and manual refinement

We retrieved the reference genome assemblies (including both the nuclear and mitochondrial genome) of *S. cerevisiae* (The *Saccharomyces* Genome Database (SGD) version of strain S288c) from SGD (<u>http://downloads.yeastgenome.org/sequence/S288C_reference/</u>) (version R64-1-1). We obtained the reference nuclear genome assembly of *S. paradoxus* (strain CBS432) from the Saccharomyces Genome Resequencing Project (SGRP) data depository (<u>ftp://ftp.sanger.ac.uk/pub/users/dmc/yeast/latest/misc/para2/ref/genome.fa</u>) and its mitochondrial genome assembly from NCBI Genbank (accession number: JQ862335).

For each polished PacBio assembly, we first used RepeatMasker (v4.0.5) to soft-mask all the repetitive regions (option: -species fungi -xsmall -gff). The soft-masked assemblies were subsequently aligned to the reference genome using the nucmer program from the MUMmer (v3.23) package (Kurtz et al. 2004) for chromosome assignment. For most chromosomes, we have a single contig covering the entire chromosome. For the cases where assembly gaps occurred in the middle of chromosomes, we performed manual gap closing by referring to the assemblies that we generated in the pilot phase of this project. The only gap that we were unable to close is the highly repetitive rDNA array (usually consisting 100-200 tandem copies) on chrXII. The *S. cerevisiae* reference genome used a 17,357 bp sequence of two tandemly arranged rDNA copies to represent this complex region. For our assemblies, we trimmed off the partially assembled rDNAs at this gap and re-linked the two contigs with 17,357 bp Ns to keep consistency.

The mitochondrial genomes of the 12 strains were recovered by single contigs in the raw HGAP assemblies. We further circularized them and reset their starting position as the *ATP6* gene using Circlator (v1.1.4) (Hunt et al. 2015).

### Illumina sequencing, reads mapping, and error correction

In addition to the PacBio sequencing, we also sequenced each strain with deep Illumina paired-end sequencing (~200X-500X) at Institut Curie (Paris, France). We examined the raw Illumina reads via FastQC (v0.11.3) and performed adapter-removing and quality-based trimming by trimmomatic (v0.33) (Bolger et al. 2014) (trimming options: ILLUMINACLIP:adapters.fa:2:30:10 SLIDINGWINDOW:5:20 MINLEN:36). For each strain, the trimmed reads were mapped to the corresponding PacBio assemblies by BWA (v0.7.12) (Li and Durbin 2009). The resulting reads alignments were subsequently processed by samtools (v1.2) (Li et al. 2009), picard tools (v1.131) (<u>http://broadinstitute.github.io/picard/</u>) and GATK (v3.5-0) (McKenna et al. 2010). The Pilon pipeline (v1.12) (Walker et al. 2014) was further used to polish the PacBio assemblies by correcting remaining sequencing errors based on the Illumina reads alignments. The Pilon-corrected assemblies were used as the final assemblies for our downstream analysis.

### Sequencing error rate evaluation for the final PacBio assemblies

Eight of our 12 strains have been previously sequenced using Illumina technology with moderate-to-high depth (Bergström et al. 2014). We retrieved the raw reads of this study and aligned them to our PacBio assemblies (both before and after Pilon correction) following the same protocol described above. The SNPs and Indels were called by FreeBayes (v1.0.1-2) (Garrison and Marth 2012) (option: -p 1) to assess the performance of Pilon correction and estimate the remaining error rate in our final assemblies. The raw SNP and Indel calls were further filtered by the vcffilter tool from vcflib (<u>https://github.com/vcflib/vcflib</u>) with the filter expression: "QUAL > 30 & QUAL / AO > 10 & SAF > 0 & SAR > 0 & RPR > 1 & RPL > 1".

### Assembly completeness comparison for different annotation features

We compared our S288c PacBio assembly with three published *S. cerevisiae* assemblies based on PacBio, Oxford Nanopore and Illumina MiSeq sequencing technologies respectively (Kim et al. 2014; Goodwin et al. 2015). We aligned these three assemblies as well as our S288c PacBio assembly to the *S. cerevisiae* reference genome using the nucmer program from the MUMmer (v3.23) package (Kurtz et al. 2004). The nucmer alignments were filtered by delta-filter (from the same package) (option: -1). We converted the output file to the “BED” format and use bedtools (v2.15.0) (Quinlan and Hall 2010) to calculate the intersection between our genome alignment and various annotation features (e.g. chromosomes, genes, retrotransposable elements, telomeres, etc) of the *S. cerevisiae* reference genome. The percent coverage of these annotation features by different assemblies was summarized accordingly.

### Centromeres annotation

For *S. cerevisiae*, centromere annotation of the reference genome is available from SGD and the corresponding sequences were retrieved as the queries. For *S. paradoxus*, the query sequences were collected from three different studies (Kellis et al. 2003; Liti et al. 2009a; Bensasson 2011), which covered 15 centromeres. All these *S. cerevisiae and S. paradoxus* centromere queries were searched against our PacBio assemblies by Exonerate (v2.2.0) (Slater and Birney 2005) (options: --showvulgar no --showcigar no --showalignment no --showtargetgff yes --bestn 1). The chrVIII centromere in *S. paradoxus* was not annotated in any of those previous studies. For this centromere, we used Dr. Bensasson’s yeast centromere annotation script (available from GitHub: <u>https://github.com/dbensasson/bensasson11</u>) to perform *de novo* annotation based on consensus sequence profile of yeast centromeres. All annotated centromeres were further verified based on their flanking genes. Several chromosomes from the strain UWOPS03-461.4, UFRJ50816 and UWOPS91-917.1 are involved in large-scale interchromosomal rearrangements. These rearranged chromosomes were numbered based on the identity of the centromeres that they are containing.

### Annotation of the protein coding genes and tRNA genes

For nuclear genes, we set up an integrative pipeline that combines three existing annotation tools to form an evidence-leveraged final annotation. First, we used the RATT package (Otto et al. 2011) to directly transfer the *S. cerevisiae* reference genome annotation to our annotated genome based on whole genome alignment. The “RATT.config_euk” file shipped with RATT was used for our transfer. The seqret program from the EMBOSS package (Rice et al. 2000) was used for format conversion between the “gff” and “embl” formats. A custom Perl script was used to further convert the seqret’s gff output file to properly formatted gff3 file. Furthermore, we used the Yeast Genome Annotation Pipeline (YGAP) online pipeline (Proux-Wéra et al. 2012) to annotate our genome assemblies with default option (except the option “ordering scaffolds by size”). YGAP is a pipeline specifically designed for annotating yeast genomes based on gene sequence homology and synteny conservation curated in the Yeast Gene Order Browser (YGOB) database. A custom Perl script was used to convert the YGAP annotation output to gff3 format while removing dubious open reading frames (ORFs) labeled by the YGAP pipeline. These dubious ORFs might be frameshifted or untranslable or overlapping with other ORFs. Also, we noticed that some ORFs annotated by YGAP might be out-of-frame (i.e. the annotated genomic coordinates went beyond the actual scaffold length) or redundant. These ORFs were further removed by our Perl script. Finally, we used the Maker pipeline (v2.31.8) (Holt and Yandell 2011) to perform *de novo* gene discovery with EST/proteome alignment support. For the Maker pipeline, repeatmasking was first performed by RepeatMasker (v4.0.5) with configuration options of “model_org=fungi” and “softmask=1”. *Ab initio* gene prediction was performed by SNAP (release 2013-11-29) (Korf 2004) and AUGUSTUS (v3.1.0) (Stanke et al. 2004) respectively with pre-trained gene prediction parameters. For SNAP, the pre-trained HMM parameter file was downloaded from GitHub (<u>https://github.com/hyphaltip/fungi-gene-prediction-params/blob/master/params/SNAP/saccharomyces_cerevisiae_S288C.hmm</u>). For AUGUSTUS, we used the parameter file for *Saccharomyces* that shipped with AUGUSTUS. In complementary to *ab initio* gene prediction, the Maker pipeline also performs EST/protein alignment to further assess the automatically predicted gene models. For the EST data, we retrieved it from FungiDB (<u>http://fungidb.org/common/downloads/release-3.2/Scerevisiae_/fasta/</u>). For protein data, we combined the proteomes from several sources: the *S. cerevisiae* proteome from SGD (<u>http://downloads.yeastgenome.org/sequence/S288C_reference/orf_protein/</u>), the protein sequences of characterized non-reference *S. cerevisiae* ORFs documented in SGD, the protein sequences of non-reference *S. cerevisiae* ORFs identified in previous studies (Novo et al. 2009; Bergström et al. 2014; Song et al. 2015), the proteomes of *S. paradoxus* (strain CBS432), *S. mikatae* (strain IFO1815), *S. kudriavzevii* (strain IFO1802), *S. kudriavzevii* (strain ZP591), and *S. bayanus var. uvarum* (strain CBS7001) based on Scannell et al. (Scannell et al. 2011), the proteome of *S. arboricolus* (strain H6) based on Liti et al. (Liti et al. 2013), and the proteome of *S. eubayanus* (strain FM1318) based on Baker et al. 2015 (Baker et al. 2015). These EST and protein sequences were aligned with the PacBio genome assemblies using blastn and blastx respectively (both from the NCBI-BLAST+ package (v2.2.30+) (Camacho et al. 2009)) and further polished by exonerate (v2.2.0) (Slater and Birney 2005). Other custom settings that we used for the Maker pipeline include: “min_contig=10000, min_protein=30, split_hit=1500, single_exon=1, single_length=250 and correct_est_fusion=1”. The gene annotations produced by RATT, YGAP, and Maker together with the EST and proteome alignment evidence generated by Maker were further leveraged by EVidenceModeler (EVM) (Haas et al. 2008) to form a final integrative version of annotation. The annotation for the *CUP1* and *ARR* cluster was further manually curated. Several genes with incomplete ORFs were also manually checked and labeled as pseudogenes if verified. In our final annotation results, ORFs overlapping with Ty retrotransposable elements, X-elements and Y’-elements were further removed. For each annotated protein-coding gene, the CDS and protein sequences were extracted using custom Perl script. We also annotated tRNA genes by tRNAscan-SE (v1.3.1) (Lowe and Eddy 1997) via the Maker pipeline.

As for mitochondrial genomes, we performed the annotation by using MFannot (<u>http://megasun.bch.umontreal.ca/cgi-bin/mfannot/mfannotInterface.pl</u>). The exon-intron boundaries of each annotated gene were manually curated based on BLAST and the 12-way whole genome alignment generated by mVISTA (Frazer et al. 2004).

### Annotation of the Ty retrotransposable elements

To systematically annotate Ty retrotransposable elements, we adopted a previously described custom library that contains *S. cerevisiae* Ty1-Ty5 and *S. paradoxus* Ty3p (Carr et al. 2012) to feed into RepeatMasker (v4.0.5) as custom library for Ty elements identification. REannotate (Pereira 2008) (v17.03.2015) was subsequently used to process the RepeatMasker output with options “-g -k <clustalw> -f <fuzzy_file1> -d 10000 -t” for Ty defragmentation. We defined the fuzzy file to treat Ty1-LTR and Ty2-LTR equivalently in the defragmentation process due to their high sequence identity and frequent recombination. The identified full-length Ty1 and Ty2 were manually curated based on their sequence alignment. We performed this Ty annotation in a two-pass manner, in which the internal sequences and LTRs of the representative full-length *S. paradoxus* Tys annotated in the first pass were further added into our Ty library before initiating the second pass. All the truncated Tys and soloLTRs were further curated based on the blastn search against our Ty library (cutoffs: identity >= 70%, aln_length >= 100 bp).

### Annotation of the core X-elements

We retrieved core X-element sequences for the *S. cerevisiae* reference genome according to the annotation from SGD and aligned them using MUSCLE (v3.8.31) (Edgar 2004). Based on the alignment, we built an HMM profile for the core X-element using the hmmbuild program (option: --dna) from the HMMER package (v3.1b2) (Eddy 1998). This HMM profile was searched against our PacBio assemblies by nhmer (from the same package) (options: -E 1e-3 --tblout) to identify the core X-element.

### Annotation of the Y’-elements

We retrieved the Y’-element sequences of the *S. cerevisiae* reference genome based on the feature annotation from SGD. There are two major classes of Y’-element for *S. cerevisiae*, the short version and the long version, differed by several large indels (Louis and Haber 1992). We aligned all the retrieved Y’-element sequences by MUSCLE (v3.8.31) (Edgar 2004). Based on this alignment, we selected the chrIX-L Y’-element as the representative query for our search. The search was performed by BLAT (Kent 2002) (option: -maxIntron=1000) with subsequent filtering by pslCDnaFilter (options: -minId=0.9 -minAlnSize=1000 -bestOverlap -filterWeirdOverlapped).

### Orthology group identification

For nuclear protein coding genes, we used proteinortho (v5.11) (Lechner et al. 2011) to identify gene orthology for all of our sequenced strains together with the well-annotated *S. cerevisiae* reference proteome from SGD (with dubious ORFs excluded) as well as the proteomes of other six *sensu stricto* species: *S. mikatae* (strain IFO1815T), *S. kudriavzevii* (strain IFO1802T), *S. kudriavzevii* (strain ZP591), *S. arboricolus* (strain H6), *S. eubayanus* (strain FM1318) and *S. bayanus var. uvarum* (strain CBS7001). The orthology identification took into account both sequence similarity and synteny conservation (the PoFF feature (Lechner et al. 2014) of proteinortho). The SGD systematic gene names were further mapped to our annotated protein coding genes according to the identified orthology.

### Phylogenetic reconstruction

For nuclear genes, we performed the phylogenetic analysis based on those one-to-one orthologs that are shared across all 18 strains (seven *S. cerevisiae* + five *S. paradoxus* + six outgroups) using two complementary approaches (the concatenated sequence tree approach and the consensus gene tree approach). For each ortholog, we used MUSCLE (v3.8.1551) (Edgar 2004) to generated protein sequence alignment and used PAL2NAL (v14) (Suyama et al. 2006) to build codon alignment based on the corresponding protein sequence alignment. For the concatenated sequence approach, we generated a concatenated codon alignment of all individual orthology groups and fed it into RAxML (v8.2.6) (Stamatakis 2014) for maximum likelihood (ML) tree building. The entire codon alignment was further partitioned by the first, second and third codon positions. The GTRGAMMA model was used for phylogenetic inference. The rapid bootstrapping method built in RAxML was used to assess the stability of internal nodes (option: -# 100 for 100 rapid bootstrap searches). The final ML tree was visualized in FigTree (v1.4.2) (http://tree.bio.ed.ac.uk/software/figtree/). For the consensus gene tree approach, we built individual gene trees with RAxML using the method as we used for the concatenated tree. Subsequently, we used ASTRAL (v4.7.12) (Mirarab et al. 2014) to estimate consensus species tree based on the topology of individual gene trees. The normalized quartet score was calculated to assess the reliability of the final species tree given individual gene trees. For mitochondrial genes, we performed the phylogenetic analysis based on the six one-to-one orthologous genes (*ATP6*, *ATP8*, *COB*, *COX1*, *COX2* and *COX3*) following the same protocol.

### Relative rate test

To test the rate heterogeneity between *S. cerevisiae* and *S. paradoxus* in molecular evolution, we constructed 3-way sequence alignment by sampling one strain for each species together with *S. mikatae* as the outgroup. The sequences were drawn from the concatenated protein alignment built from one-to-one orthologs in the nuclear genome. The extracted sequences were fed into MEGA (v6.06-mac) (Tamura et al. 2013) for Tajima’s relative rate test (Tajima 1993). We conducted this test for all possible *S. cerevisiae* versus *S. paradoxus* strain pairs.

### Molecular dating

Since no yeast fossil records can be used for reliable calibration, we performed the molecular dating analysis based on a relative time scale. We used the phylogenetic tree constructed from one-to-one orthologs in the nuclear genome as the input and performed least-square based fast dating with LSD (To et al. 2016) (options: -c -v -s). We specified *S. bayanus var. uvarum* CBS7001 and *S. eubayanus* FM1318 as outgroups for this analysis. Based on the resulting chronogram, we further summed up the lengths of all branches in the corresponding species-specific clade as the cumulative diversification time for the *S. cerevisiae* and *S. paradoxus* strains respectively.

### Conserved synteny blocks identification

We used the SynChro program from the CHROnicle package (version: January 2015) (Drillon et al. 2013, 2014) to identify conserved synteny blocks. We prepared the input files for SynChro with custom Perl scripts to provide information about various annotated features (centromere, protein-coding genes, tRNAs, and Tys) together with the genome assembly and proteome sequences. SynChro subsequently performed all possible pairwise comparisons to identify synteny blocks shared in the given strain pair. Multiple plots were also generated by SynChro for easy visualization of the identified synteny blocks.

### Subtelomere definition and chromosome partitioning

Yeast subtelomeres are known for a few general properties such as low gene density, low synteny conservation, and silent chromatin state. An often-used definition is 20-30 kb from the chromosome-ends. However, this definition seems arbitrary in a sense that it treats all subtelomeres indiscriminately. In this study, we defined yeast subtelomeres based on the change of gene synteny conservation profile across the 12 strains. For each chromosome arm, we examined the syntenic blocks shared across all the 12 strains and used the most distal syntenic blocks to define the distal boundary for the chromosomal cores (Table S11). In parallel, we defined the proximal boundary of the chromosome-end region for this chromosome arm based on the first occurrence of yeast telomere associated sequences (i.e. core X-and Y’-element). The region between these two boundaries was defined as the corresponding subtelomere with 400 bp interstitial transition zones on both sides (Figure S3). Each defined subtelomere was named according to the ancestral chromosome identity of the core region that it attaches to.

### Identification of balanced and unbalanced structural rearrangements

We identified balanced genome rearrangements (inversion, translocation, and transposition) by the ReChro from the CHROnicle package (version: January 2015) (Drillon et al. 2013, 2014). We set the synteny block stringency parameter delta=1 for the main analysis. A complementary run was performed with delta=0 to identify single gene inversions. As for unbalanced genome rearrangements (large insertion, deletion and duplication), we first generated whole genome alignment for every strain pair by nucmer (Kurtz et al. 2004) (options: -maxmatch -c 500) and submitted the results to the Assemblytics web server (http://assemblytics.com/) (Nattestad and Schatz 2016) for identifying all potential insertions, deletions and duplications/contractions in the corresponding pairwise comparison. All candidates identified by Assemblytics were further compared with our gene annotation by bedtools intersect (Quinlan and Hall 2010) (options: -wo -F 0.9) to only keep those candidates that are overlapping with protein coding genes. All post-filtered balanced and unbalanced structural rearrangements were manually checked with chromosome-scale dotplots using Gepard (v1.30) (Krumsiek et al. 2007) for final verification. Here we only focused on those structural rearrangements occurred in core regions since the rampant ectopic subtelomeric reshuffling would introduce considerable noise into our synteny-based detection. All verified core-region rearrangements were mapped to the phylogeny of the 12 strains based on the maximum parsimony principle.

### Gene ontology analysis

The CDS sequences of all SGD reference genes (with dubious genes excluded) were BLAST against NCBI non-redundant (nr) database using blastx (E-value cutoff = 1E-3) and further annotated by BLAST2GO (v.3.2) (Conesa et al. 2005; Götz et al. 2008) to generate gene ontology (GO) mapping for the genome-wide background gene set. For all the genes involved in unbalanced structural rearrangements, their corresponding orthologs in the SGD reference gene set (when available) was used to compile a non-redundant test gene set. We performed Fisher’s exact test (Fisher 1922) to detect significantly enriched GO terms of our test gene set relative to the genome-wide background. False discovery rate (FDR) (cutoff: 0.05) (Benjamini and Hochberg 1995) was used for multiple correction.

### Molecular evolutionary rate and CNV estimation

For the one-to-one orthologs in each strain pair, we calculate synonymous substitution rate (dS), nonsynonymous substitution rate (dN) and nonsynonymous-to-synonymous substitution rate ratio (dN/dS) using the yn00 program from the PAML package (v4.8a) (Yang 2007). Yang & Nielsen (2000) model (Yang and Nielsen 2000) was used for this calculation. As for measuring copy number variation (CNV), we calculated the proportion of genes involved in CNV (i.e. those are not one-to-one orthologs) in the two compared strains. We denoted this measurement as *P_CNV_*, a quantity analogous to the *P*-distance in sequence comparison. The Poisson distance correction was further applied to account for multiple changes at the same gene loci. The Poisson corrected distance *D_CNV_* can be given as *D_CNV_* = −ln (1 − *R_CNV_*). Then the CNV accumulation rate (*R_CNV_*) can be calculated as *D_CNV_* = *R_CNV_*/*2T*, in which *T* is the diversification time of the two compared strains obtained from our molecular dating analysis. The calculation values for dN/dS, CNV proportion, and CNV accumulation rate were further summarized by “core genes” and “subtelomeric genes” based on our genome partitioning described above.

### Subtelomeric homology search

For each defined subtelomeric region, we hard masked all the Ty-related features (full-length Ty, truncated Ty and Ty soloLTRs) involved and then used the masked sequence to search against all the other subtelomeric regions to detect shared sequence homology. The search was performed by BLAT (Kent 2002) (options: -noHead -stepSize=5 -repMatch=2253 -minIdentity=80 -t=dna -q=dna -mask=lower -qMask=lower). We further used pslCDnaFilter (options: -minId=0.9 -minAlnSize=1000 -bestOverlap -filterWeirdOverlapped) to filter out trivial signals and used pslScore to calculate sequence alignment score for those filtered BLAT matches. Since the alignment scores for a given subtelomere pair are not exactly symmetrical, we considered the average score between the two ordered pairs in such cases. Such subtelomeric homology search was carried out for both within-strain and cross-strain comparison and subtelomere pairs with strong homology (BLAT alignment score >= 5000 and sequence identity >= 90%) were considered.

### Hierarchical clustering analysis and reshuffling rate calculation for orthologous subtelomeres

For all the strains within the same species, we performed pairwise comparisons of their subtelomeric regions to identify conserved orthologous subtelomeres in any given strain pairs based on homology search described above. For each strain pair, the proportion of conserved orthologous subtelomeres was calculated as a measurement of the overall subtelomere conservation between the two compared strains. Such measurements were converted into a distance matrix, based on which the hierarchical clustering analysis was further performed by R (v3.1) (R Developement Core Team 2015). We measured the reshuffling rate of orthologous subtelomeres (*R_reshuffling_*) similarly to how we calculated the CNV accumulation rate (*R_CNV_*). For any given strain pair, we first calculated the proportion of the non-conserved orthologous subtelomeres in this strain pair as *P_reshuffling_*, then the subtelomeric reshuffling rate *R_reshuffling_* can be calculated as *R_reshuffling_* = −ln (1 − *P_reshuffling_*)/*2T*, in which *T* is the diversification time of the two compared strains.

### Phenotyping the growth rates of yeast strains in copper-and arsenite-rich medium

The homozygous diploid versions of the 12 strains were pre-cultured in Synthetic Complete (SC) medium for overnight to saturation. To examine their conditional growth rates in copper-and arsenite-rich environment, we mixed 350 µl conditional media (CuCl_2_ (0.38 mM) and arsenite (As[III], 3 mM) for the two environment respectively) with 10 µl saturated culture to the wells of Honeycomb plates (9502550, Bioscreen). Oxygen permeable films (Breathe-easy, BEM-1, Diversified Biotech) were placed on top of the plates to enable a uniform oxygen distribution throughout the plate. The automatic screening was done with Bioscreen analyser C (Thermic Labsystems Oy, Finland) at 30°C for 72 hours, measuring in 20 minute intervals using a wide-band filter at 420-580 nm (Warringer and Blomberg 2003). Growth data pre-processing and phenotypic trait extraction was performed by PRECOG (Fernandez-Ricaud et al. 2016).

### Linkage analysis in diploid *S. cerevisiae* hybrids

A total of 826 phased outbred lines (POLs) were constructed and phenotyped in the same fashion as previously described (Hallin et al. 2016). Briefly, the strains YPS128 and DBVPG6044 were mated and advanced intercrossed lines were created by successive sporulation and crossing. The resulting haploid advanced intercrossed lines were sequenced and further crossed in a number of combinations to yield the 826 POLs used for the analysis. The POL diploid genotypes can be accurately inferred from the haploid advanced intercrossed lines. Phenotyping of the POLs, each with four replicates, was performed using Scan-o-Matic (Zackrisson et al. 2016) on solid agar plates (0.14% Yeast Nitrogen Base, 0.5% ammonium sulphate, 2% (w/v) glucose and pH buffered to 5.8 with 1% (w/v) succinic acid, 0.077% Complete Supplement Mixture (CSM, Formedium™), 2% agar) supplemented with one out of four different arsenite concentrations (0, 1, 2, and 3mM). Using the deviations between the POLs and their estimated parental as phenotypes to combat population structure issues (Hallin et al. 2016), QTLs were mapped using the scanone() function in R/qtl (Broman et al. 2003) with the marker regression method.

## Acknowledgements

We thank Guenola Drillon for the help with using the program CHROnicle. We thank Olivier Croce and Roberto Marangoni for the help with maintaining the computing server and various bioinformatics tools. We thank Liti lab technician Agnès Llored for preparing some of strain and DNA samples. This work was supported by ATIP-Avenir (CNRS/INSERM), Fondation ARC pour la Recherche sur le Cancer (grant number PJA20151203273), European Commission (Marie Curie Career Integration Grants (FP7-PEOPLE-2012-CIG) grant number 322035), Agence Nationale de la Recherche (grant number ANR-13-BSV6-0006-01 and Labex SIGNALIFE ANR-11-LABX-0028-01), Cancéropôle PACA (AAP émergence 2015) and DuPont Young Professor Award to GL, Wellcome Trust (grant number WT098051) to RD, and Vetenskapsrådet (The Swedish Research Council) (grant number 325-2014-4605) to JW. JXY is supported by a postdoctoral fellowship from Fondation ARC pour la Recherche sur le Cancer (grant number n°PDF20150602803). JL is supported by a postdoctoral fellowship from Fondation ARC pour la Recherche sur le Cancer (grant number n°PDF20140601375). JH is supported by the Labex SIGNALIFE program from Agence Nationale de la Recherche (grant number ANR-11-LABX-0028-01).

## Additional information

### Author contributions

J-XY, conceived, designed, and performed the bioinformatics analysis, wrote the manuscript.

JL, prepared DNA samples for sequencing, performed the experiment on verifying structural rearrangement, contributed to the manuscript.

LA, performed the PacBio sequencing and helped with diagnosing the assembly pipeline.

JH, performed experiments and data analysis for phenotyping, contributed to the manuscript.

KP, performed experiments and data analysis for phenotyping, contributed to the manuscript.

KO, performed the PacBio sequencing and ran the standard assembly pipeline.

AB, helped with discussion on data analysis and manuscript preparation.

PC, performed the PacBio sequencing for the pilot phase project.

JW, designed the phenotyping experiment and helped with data interpretation.

MCL, helped with the analysis on measuring the sequence homology for subtelomeres.

GF, helped with discussion on data analysis and manuscript preparation.

RD, conceived and designed the study.

GL, conceived, designed, and guided the study, wrote the manuscript.

## Data access

### The following datasets were generated

Yue JX, Li J, Aigrain L, Hallin J, Persson K, Oliver K, Bergström A, Coupland P, Warringer J, Consentino Lagomarsino M, Fischer G, Durbin R and Liti G, 2016.

### The PacBio sequencing for 12 representative strains from *Saccharomyces cerevisiae* and its wild relative *Saccharomyces paradoxus*

<u>http://www.ebi.ac.uk/ena/data/view/PRJEB7245</u>

Publicly available at the EBI European Nucleotide Archive (accession no: PRJEB7245).

Yue JX, Li J, Aigrain L, Hallin J, Persson K, Oliver K, Bergström A, Coupland P, Warringer J, Consentino Lagomarsino M, Fischer G, Durbin R and Liti G, 2016.

### The Illumina sequencing for 12 representative strains from *Saccharomyces cerevisiae* and its wild relative *Saccharomyces paradoxus*

<u>http://www.ncbi.nlm.nih.gov/bioproject/340312</u>

Publicly available at the NCBI Short Reads Archive (accession no: PRJNA340312).

Yue JX, Li J, Aigrain L, Hallin J, Persson K, Oliver K, Bergström A, Coupland P, Warringer J, Consentino Lagomarsino M, Fischer G, Durbin R and Liti G, 2016.

### The final assemblies, annotations and extracted CDS sequences, and protein sequences for 12 representative strains from *Saccharomyces cerevisiae* and its wild relative *Saccharomyces paradoxus*

<u>https://yjx1217.github.io/Yeast_PacBio_2016/data/</u>

Publicly available at our dedicated website for this project hosted via GitHub.

### The following previously published datasets were used

Liti G, Carter DM, Moses AM, Warringer J, Parts L, James SA, Davey RP, Roberts IN, Burt A, Koufopanou V, Tsai IJ, Bergman CM, Bensasson D, O'Kelly MJT, van Oudenaarden A, Barton DBH, Bailes E, Nguyen AN, Jones M, Quail MA, Goodhead I, Sims S, Smith F, Blomberg A, Durbin R and Louis EJ, 2009

### Population genomics of domestic and wild yeasts

<u>ftp://ftp.sanger.ac.uk/pub/users/dmc/yeast/latest/</u>

Publicly available at the dedicated FTP directory for the SGRP project at Sanger Institute.

Bergström A, Simpson JT, Salinas F, Barré B, Parts L, Zia A, Nguyen Ba AN, Moses AM, Louis EJ, Mustonen V, Warringer J, Durbin R, and Gianni Liti G, 2014

### A High-Definition View of Functional Genetic Variation from Natural Yeast Genomes

<u>ftp://ftp.sanger.ac.uk/pub/users/dmc/yeast/SGRP2/input/strains/</u>

Publicly available at the dedicated FTP directory for the SGRP2 project at Sanger Institute.

Kim KE, Peluso P, Babayan P, Yeadon PJ, Yu C, Fisher WW, Chin CS, Rapicavoli NA, Rank DR, Li J, Catcheside DE, Celniker SE, Phillippy AM, Bergman CM, Landolin JM, 2014

### Long-read, whole-genome shotgun sequence data for five model organisms

<u>http://www.ncbi.nlm.nih.gov/assembly/GCA_000773925.1</u>

Publicly available at the NCBI Genbank (accession no: ASM77392v1).

Goodwin S, Gurtowski J, Ethe-Sayers S, Deshpande P, Schatz MC, McCombie WR.

### Oxford Nanopore sequencing, hybrid error correction, and de novo assembly of a eukaryotic genome

<u>http://schatzlab.cshl.edu/data/nanocorr/</u>

Publicly available via a dedicated webpage hosted at the Schatz lab.

---

**Figure S1.**
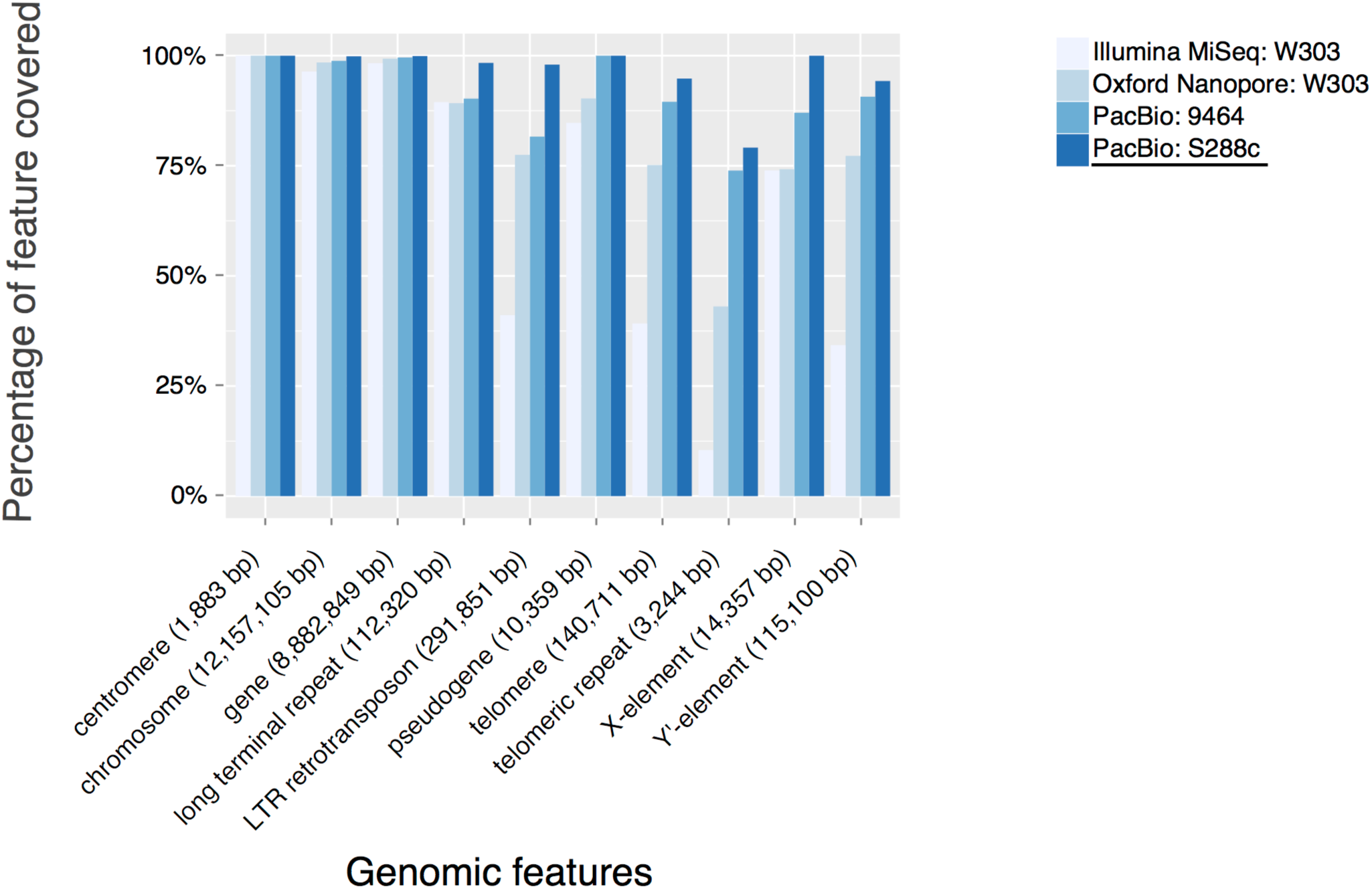
**Percentage of genomic features annotated in the *S. cerevisiae* reference genome covered by different *S. cerevisiae* assemblies.** Our PacBio assembly (strain S288c, underscored) shows all-around best completeness when compared with other *S. cerevisiae* assemblies from previous studies (Kim et al. 2014; Goodwin et al. 2015) using Illumina MiSeq (strain W303), PacBio (strain 9464), and Oxford Nanopore (strain W303) technologies. The strain W303 and 9464 sequenced by previous studies are phylogenetically very close to S288c. The difference in assembly completeness is especially pronounced for repetitive features such as those related to telomeres.

**Figure S2.**
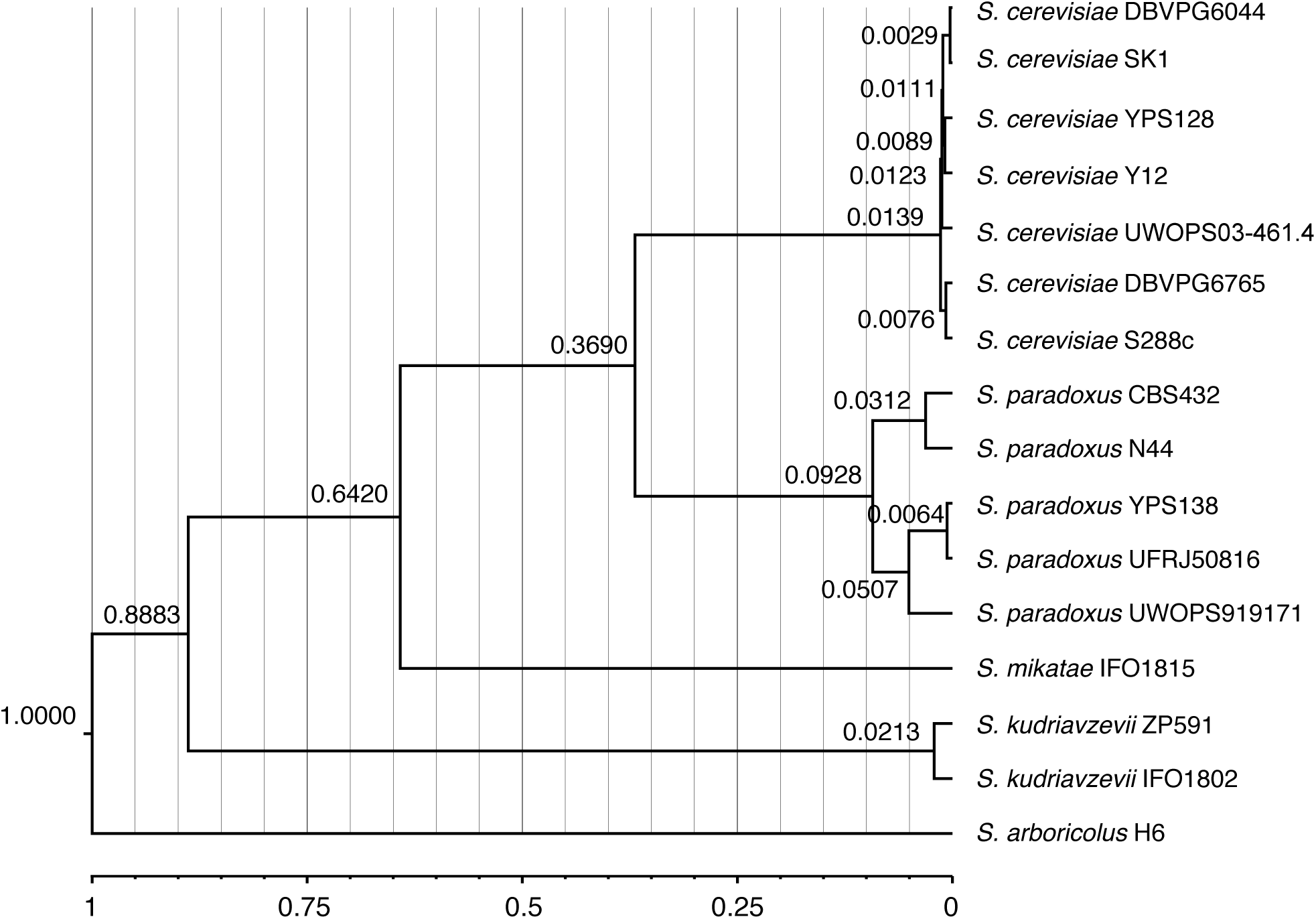
**Molecular dating for the evolutionary history of the *Saccharomyces sensu stricto* yeasts.** The number at each internal node denotes the corresponding diversification or divergence time in relative measurement. The scale at the bottom denotes the relative time frame. The *S. bayanus var. uvarum* strain CBS7001 and *S. eubayanus* strain FM1318 were not included in this plot because they were used as outgroups for this analysis.

**Figure S3.**
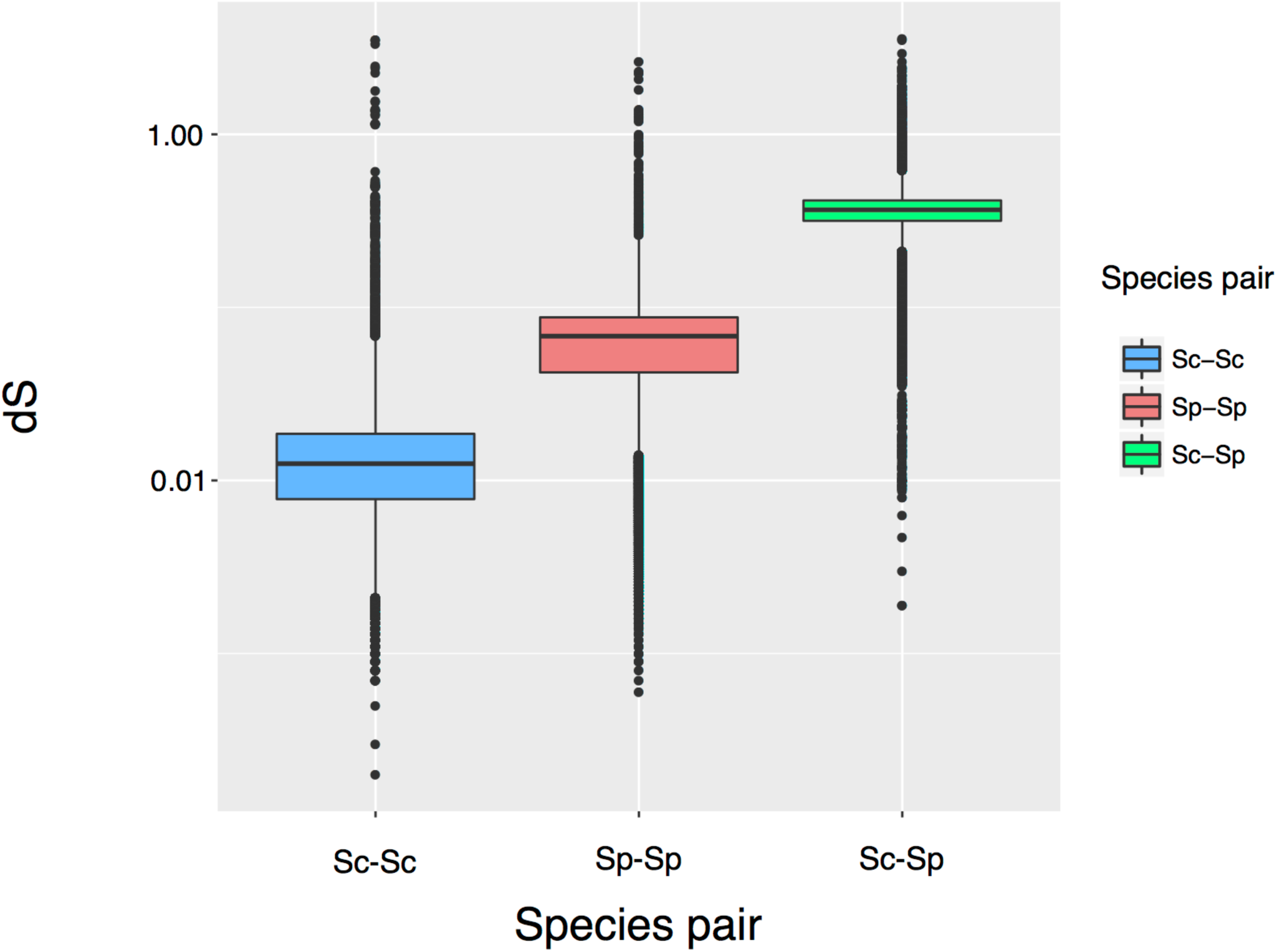
**Synonymous substitution rates (dS) in both within-and cross-species comparisons.** Three comparison scales were examined: within *S. cerevisiae* (Sc-Sc), within *S. paradoxus* (Sp-Sp) and between the two species (Sc-Sp).

**Figure S4.**
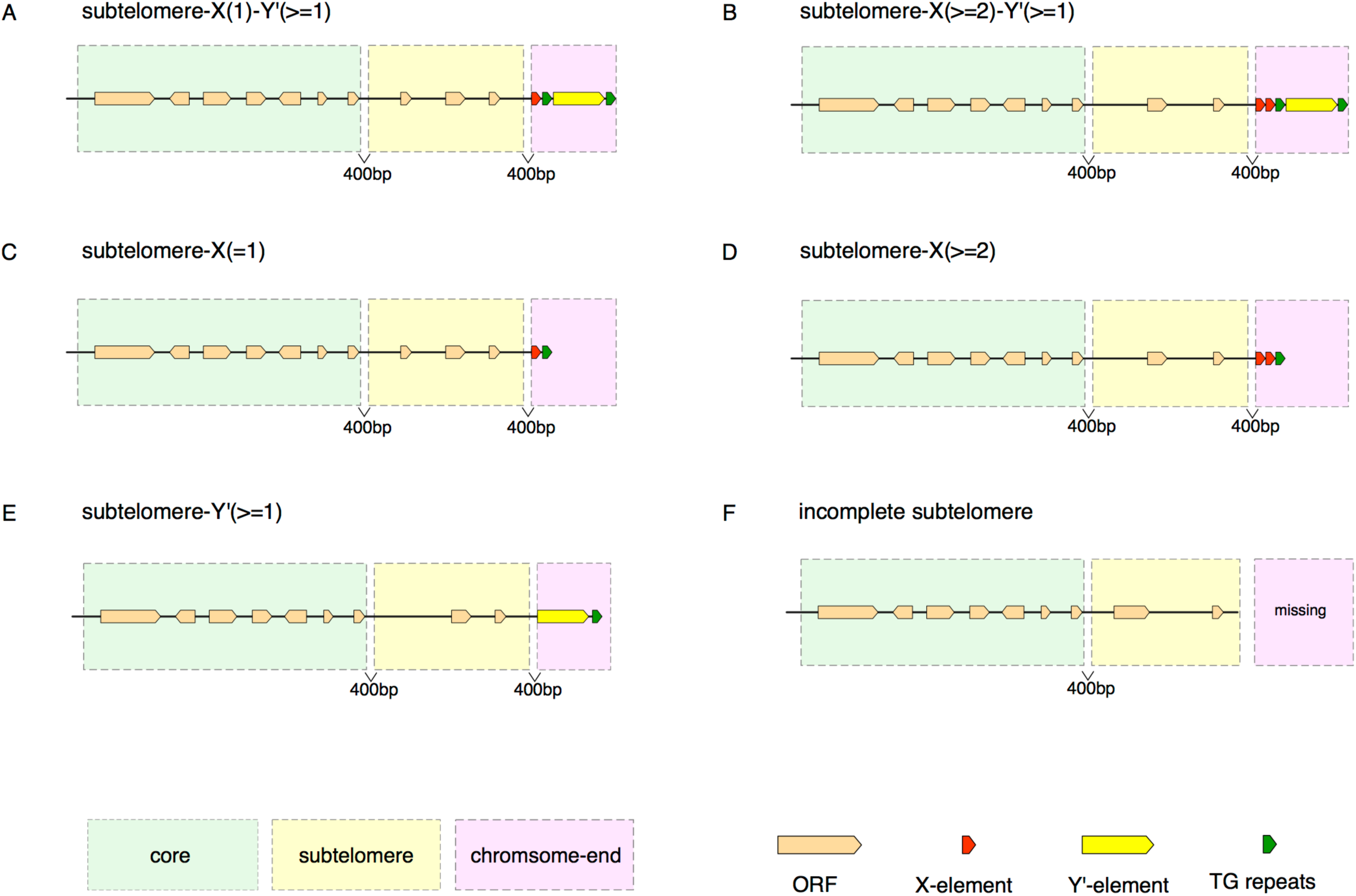
**Natural variation of chromosome-end structures.** The chromosome-end structures list here are based on Table S13. Numbers in parenthesis denotes the copy number of X-or Y’-elements that the corresponding chromosome-end structure could have.

**Figure S5.**
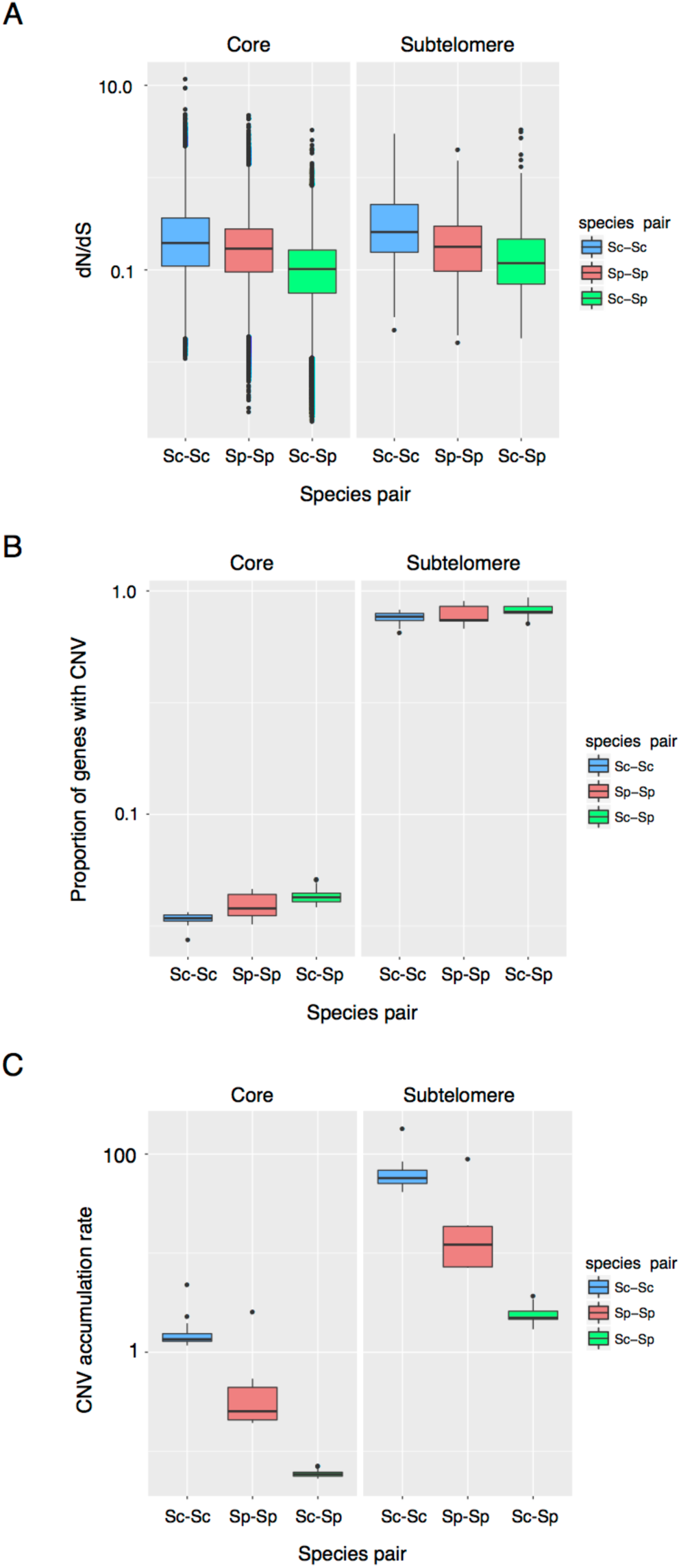
**Pronounced contrasts between cores and subtelomeres in evolutionary dynamics.** Nonsynonymous to synonymous substitution rate ratios (dN/dS), proportions of genes involved in copy number variation (CNV), and accumulation rates of CNV in both cores and subtelomeres were shown in A-C. Three comparison scales were examined: within *S. cerevisiae* (Sc-Sc), within *S. paradoxus* (Sp-Sp) and between the two species (Sc-Sp). The y-axes for proportions of genes involved in CNV (B) and accumulation rates of CNV (C) are in log-10 scales.

**Figure S6.**
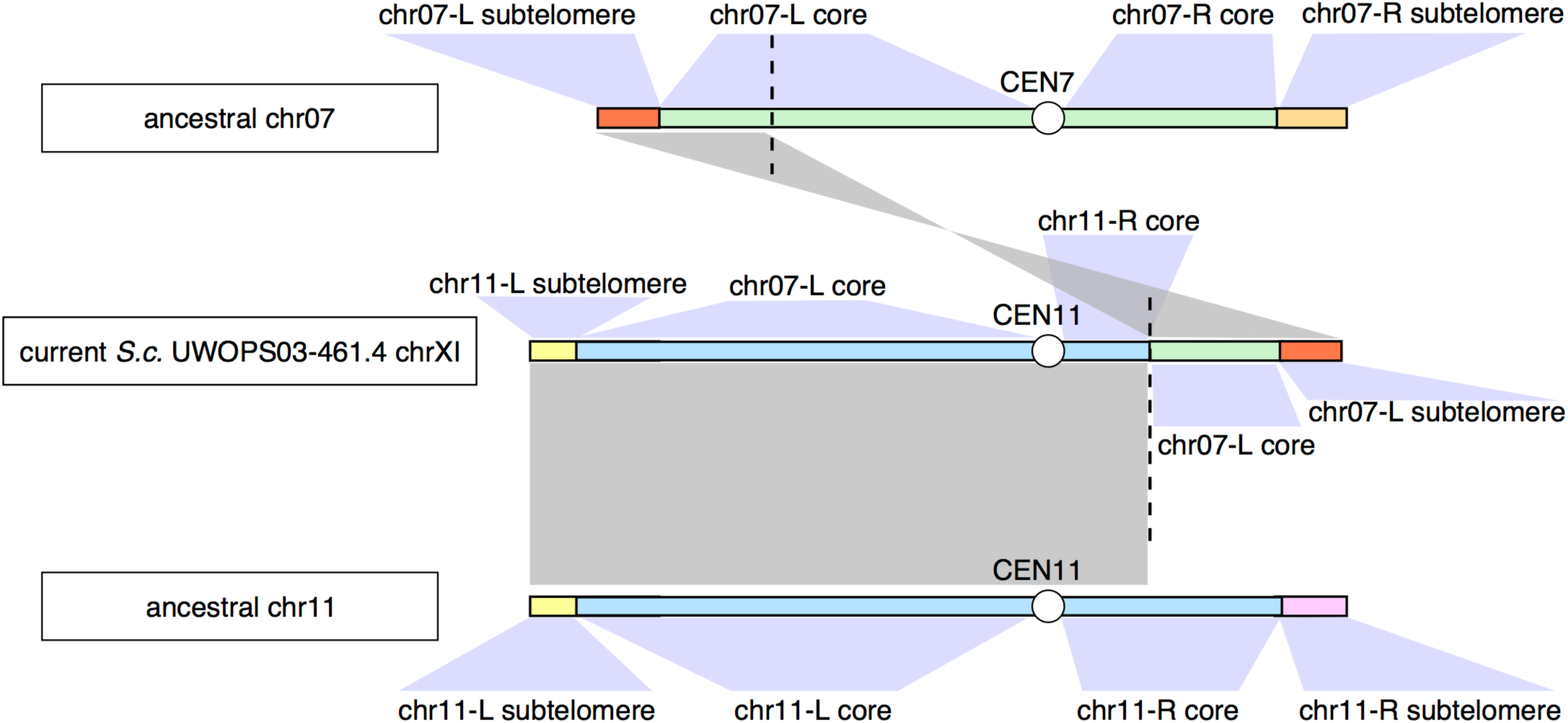
**Naming rationale of the subtelomeric region based on its ancestral identity to account for large interchromosomal rearrangements occurred in the Malaysian *S. cerevisiae* UWOPS03-461.4 and the South American and Hawaiian *S. paradoxus* (UFRJ50816 and UWOPS91-917.1 respectively).** In this example, the current chrXI (named based its centromere) of UWOPS03-461.4 contains material from both ancestral chr07 and chr11 due to a large interchromosomal rearrangement. We used grey blocks to denote the homologous relationship of different section of the current UWOPS03-461.4 chrXI relative to the ancestral chr07 and chr11.

**Figure S7.**
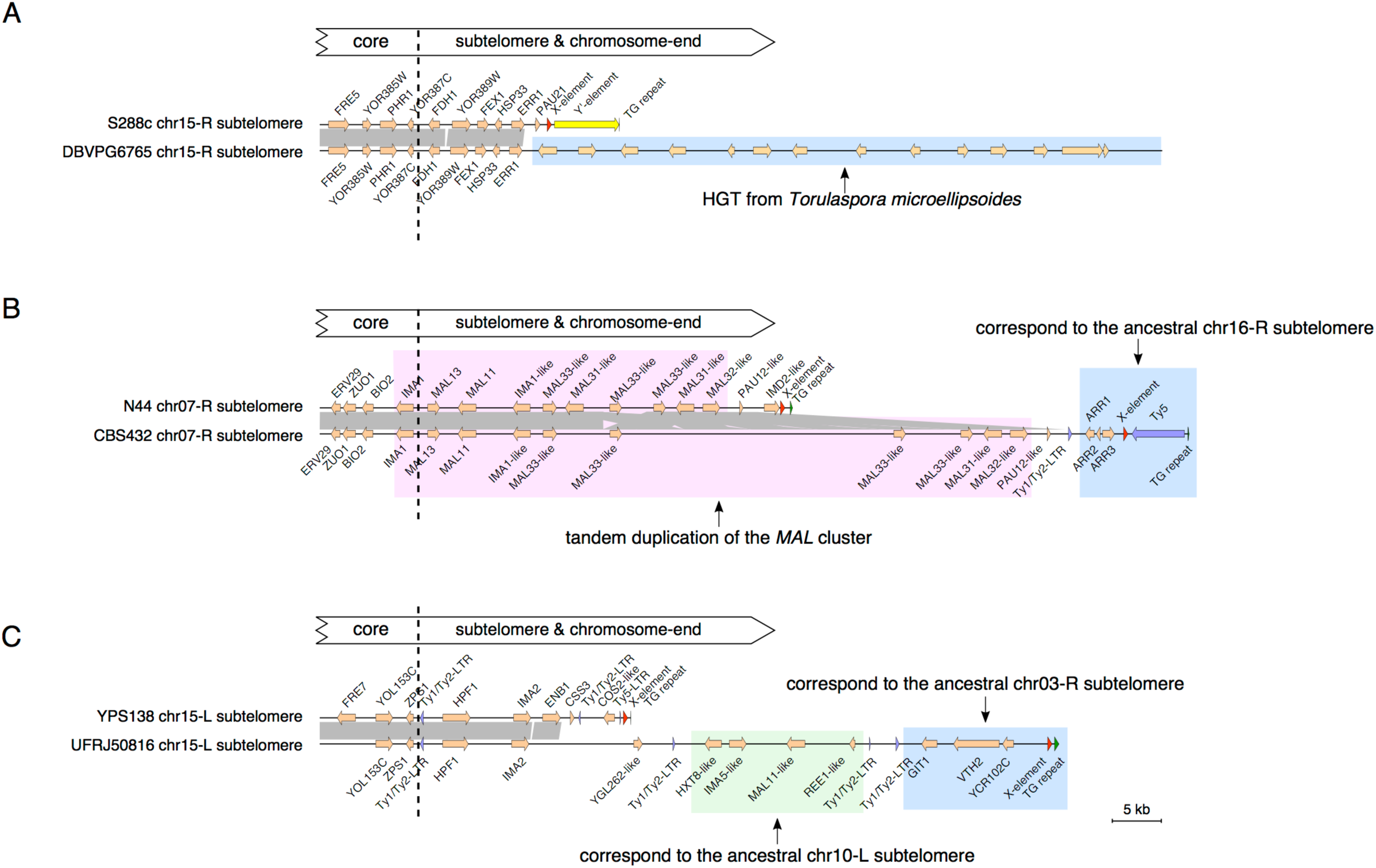
**Detailed gene maps explaining the three cases of subtelomere size expansion.** These three cases (A, B and C) correspond to Figure 4B, 4C, and 4D respectively.

**Figure S8.**
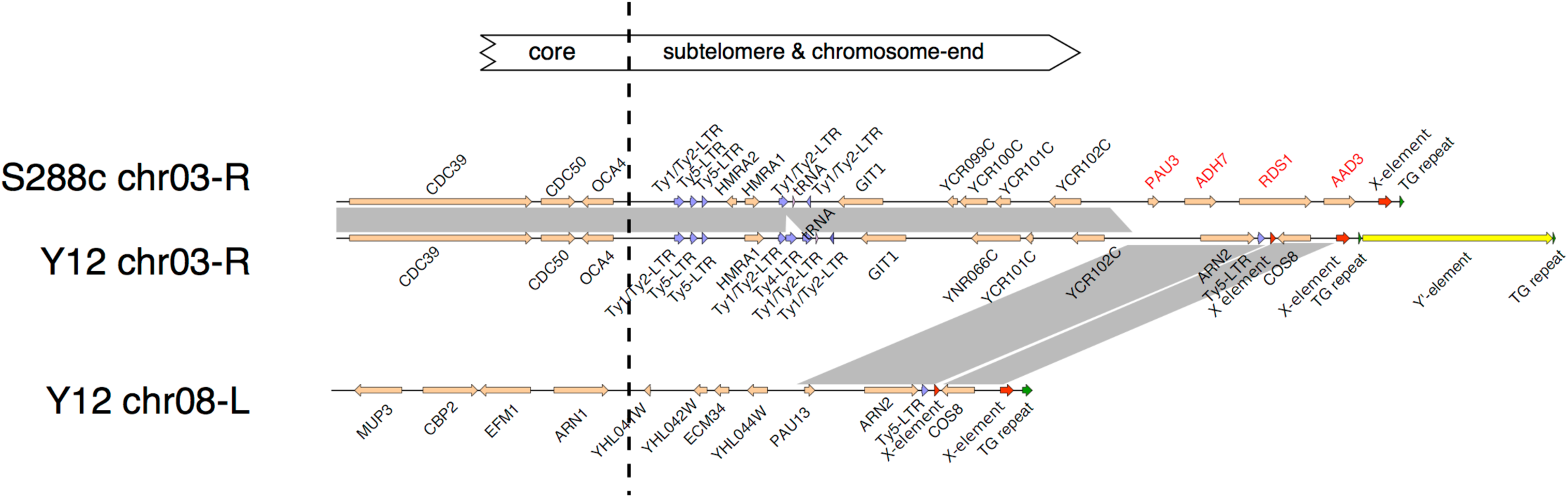
**Gene loss due to a subtelomeric duplication event in the chr03-R subtelomere of the *S. cerevisiae* strain Y12.** Four genes (*PAU3*, *ADH7*, *RDS1*, and *AAD3*) (highlighted in red) in the ancestral chr03-R (displayed here based on S288c) were lost in Y12 due to the subtelomeric duplication event from chr08-L subtelomere. Homologous regions with sequence identity >90% were highlighted in grey blocks.

## Titles for supplementary tables

Table S1: Strain sampling for this study.

Table S2: PacBio and Illumina sequencing depth.

Table S3: Pilon correction for nuclear genome assemblies.

Table S4: Pilon correction for mitochondrial genome assemblies.

Table S5: Total occurrences of different genomic features annotated in the nuclear genome.

Table S6: Total occurrences of Ty-related genomic features annotated in the nuclear genome.

Table S7: Total occurrences of different genomic features annotated in the mitochondrial genome.

Table S8. Cumulative length for different genomic features annotated in the nuclear genome.

Table S9: Cumulative length for annotated features in the mitochondrial genome.

Table S10: Distribution of group I and group II introns in the mitochondrial genome.

Table S11: The core region boundaries defined based on gene synteny conservation across the 12 strains.

Table S12: Subtelomeres involved in large-scale chromosomal rearrangements based on their ancestral locations.

Table S13: Chromosome-end structure characterized for the 12 strains.

Table S14: Enriched Gene Ontology (GO) terms of genes involved in unbalanced rearrangement.

Table S15: Pairs of subtelomeric duplication blocks within each strain.

## Titles for supplementary data

Supplementary data 1: Genomic coordinates for subtelomeres identified in the 12 strains.

Supplementary data 2: Genomic features within the identified subtelomeres.

Supplementary data 3: Balanced rearrangement events identified in the 12 strains.

Supplementary data 4: Unbalanced rearrangement events identified in the 12 strains.

Supplementary data 5: Subtelomeric duplication blocks identified in the 12 strains.

Supplementary data 6: Strain-sharing pattern of duplicated subtelomere pairs.

